# Mapping Hand Function with Simultaneous Brain-Spinal Cord Functional MRI

**DOI:** 10.1101/2025.02.27.640504

**Authors:** Valeria Oliva, Sandrine Bédard, Merve Kaptan, Dario Pfyffer, Brett Chy, Susanna Aufrichtig, Nazrawit Berhe, Akshay S. Chaudhari, Suzanne Tharin, Serena S. Hu, John Ratliff, Zachary A. Smith, Andrew C. Smith, Gary H. Glover, Sean Mackey, Christine S. W. Law, Kenneth A. Weber

## Abstract

**INTRODUCTION:** Hand motor control depends on intricate brain-spinal cord interactions that regulate muscle activity. Hand function can be disrupted by injury to the brain, spinal cord, and peripheral nerves leading to weakness and impaired coordination. Functional MRI (fMRI) can map motor-related neural activity and potentially characterize the mechanisms underlying hand weakness and diminished coordination. Although brain motor control has been extensively studied, spinal cord mechanisms remain less explored. Here we use simultaneous brain-spinal cord fMRI to map neural activity related to hand strength and dexterity across the central nervous system using force matching and finger tapping tasks. This study pioneers the use of simultaneous brain-spinal cord fMRI to comprehensively map hand function, offering novel insights into coordinated motor processing across the central nervous system.

**METHODS:** We performed simultaneous brain-spinal cord fMRI in 28 right-handed healthy volunteers (age: 40.0 ± 13.8 years, 14 females, 14 males) using a 3T GE SIGNA Premier scanner equipped with a 21-channel head-neck coil. Participants performed a force-matching task at 10%, 20%, and 30% of maximum voluntary contraction using a hand dynamometer. For the finger tapping task, participants completed button-presses at 1 Hz with a 5-button response pad for three task levels: single-digit response with the second digit only (low), single-digit response with all digits in a sequential order (medium), single-digit response with all digits in a random order (high). Visual cues and feedback were provided during the tasks.

Brain and spinal cord images were processed separately using FSL and the Spinal Cord Toolbox, with motion correction, physiological noise filtering, and spatial normalization to standard templates. Subject level activity maps were generated and entered into group level analyses to explore both activations and deactivations. For the brain, we used a mixed effect design with a voxelwise threshold of Z score > 3.10 and cluster threshold of p < 0.05. For the spinal cord, we used a fixed effect design with a voxelwise threshold of Z score > 1.64 and cluster threshold of p < 0.05. Region of interest (ROI) analyses were conducted to examine localized changes in activation across task levels

**RESULTS:** Both tasks elicited activation in motor and sensory regions of the brain and spinal cord, with graded responses in the left primary motor (M1), left primary sensory (S1) cortex, and right spinal cord gray matter across task levels. Deactivation of the right M1 and S1 was also present for both tasks. Deactivation of the left spinal cord gray matter was present in the high task level of the force matching task. The ROI analysis findings complemented the group level activity maps.

**DISCUSSION:** Our study provides a detailed map of brain-spinal cord interactions in hand function, revealing graded neural activation and inhibition patterns across motor and sensory regions. Interhemispheric inhibition, reflected in right M1 deactivation, likely restricts extraneous motor output during unilateral tasks. For force matching, the deactivation of the left ventral and dorsal horns of the spinal cord, provides the first evidence that the inhibition of motor areas during a unilateral motor task extends to the spinal cord. Whether this inhibition results from direct descending modulation from the brain or interneuronal inhibition in the cord remains to be interrogated. These findings expand our understanding of central motor control mechanisms and could inform rehabilitation strategies for individuals with motor impairments.

**CONCLUSIONS:** Our simultaneous brain-spinal cord fMRI approach provides novel insights into the neural coordination of hand function, enhancing our understanding of motor control and its modulation. This approach may offer a foundation for studying motor dysfunction in conditions such as stroke, spinal cord injury, and neurodegenerative diseases.

## 1. Introduction

Hand function, ranging from gross grasping to precision movements, depends on coordinated interactions between the brain, spinal cord, and peripheral muscles. Cortical (e.g., primary motor (M1), primary sensory (S1), SMA, premotor cortices) and subcortical (e.g., basal ganglia, cerebellum) structures contribute to planning and executing voluntary movements. These brain regions transform a movement plan from real-world coordinates to muscle-specific motor commands which descend the spinal cord and recruit motor neurons leading to the appropriate muscle contractions (Castiello & Begliomini, 2008; Witt et al., 2008). Despite extensive research on brain mechanisms of hand function, it is necessary to better understand how spinal cord activity integrates with brain motor control. Injuries or dysfunction at these levels can cause weakness and impaired coordination, yet distinguishing between cortical, subcortical, and spinal contributions remains a challenge. Simultaneous brain-spinal cord fMRI offers a unique opportunity to resolve these gaps by mapping functional interactions across the neuraxis.

Functional MRI (fMRI) has revolutionized our understanding of brain motor control, revealing cortical reorganization in stroke and Parkinson’s disease and informing rehabilitation strategies (Binkofski et al., 1999; Filippi et al., 2019; Grefkes & Fink, 2011; Hou et al., 2016; Johansen-Berg et al., 2002; Leskinen et al., 2024). Recent advances—such as improved shimming, reduced field-of-view imaging, and specialized spinal cord imaging techniques—now enable fMRI to probe spinal cord function, bridging the gap between brain and spinal motor research (Finsterbusch et al., 2012; Fratini et al., 2014; Giove et al., 2004; Islam et al., 2019; Kaptan et al., 2024). The development of simultaneous brain-spinal cord fMRI now allows for comprehensive mapping of motor function across the central nervous system (Vahdat et al., 2015). While prior studies have separately examined cortical and spinal motor activity, a unified view of how the brain and spinal cord interact during voluntary movement remains elusive (Giulietti et al., 2008; Hemmerling et al., 2023; Hernandez-Charpak et al., 2025; Kinany et al., 2023; Maieron et al., 2007; Stroman & Ryner, 2001; Weber et al., 2016). This study leverages simultaneous fMRI to elucidate these interactions, providing novel insights into the neural basis of hand strength and dexterity.

With this in mind, our study applies simultaneous brain-spinal cord fMRI to elucidate neural mechanisms underlying hand strength and dexterity. Using force-matching (i.e., hand strength) and finger-tapping (i.e., dexterity) tasks at three difficulty levels, we aim to map graded motor and sensory activity across the neuraxis. We hypothesize that (1) task difficulty will modulate activity in both brain and spinal cord regions, (2) force generation will preferentially engage primary motor and sensory cortices, and (3) interhemispheric inhibition and spinal cord deactivations will regulate unilateral motor output. By integrating brain and spinal cord imaging, this study advances our understanding of motor control and its clinical applications. Our findings may refine sensorimotor models and inform rehabilitation strategies for stroke, spinal cord injury, and neurodegenerative disorders.

## 2. Methods

### 2.1. Participants

Twenty-eight right-handed healthy volunteers (age = 40.0 ± 13.8 years, 14 females, 14 males) were recruited. Exclusion criteria included any contraindications to MRI, pain conditions, previous spine surgery, and major medical, psychiatric, and neurological conditions. Eligible volunteers received an overview of the study protocol, and then gave written informed consent. The study was approved by Stanford University’s Institutional Review Board.

### 2.2. Hand Function

We assessed handedness using the Edinburgh Handedness Inventory Short Form (Veale, 2014). We measured right hand grip strength (Jamar plus dynamometer, Performance Health Supply, Inc., Cedarburg, WI, USA) and dexterity (Jamar 9 Hole Peg Test kit, Performance Health Supply, Inc., Cedarburg, WI, USA) using the normative scores from the NIH Toolbox motor battery, which account for age, sex, education level, race, and ethnicity (Reuben et al., 2013).) Upper extremity physical function was also assessed using the Quick DASH outcome measure (Beaton et al., 2005) and PROMIS Upper Extremity Short Form 7a (version = 2.1) (Kaat et al., 2019).

### 2.3. Functional MRI Experiments

For the force matching task, participants gripped an MRI-compatible dynamometer (BIOPAC Systems, Inc., Goleta, CA, USA) with their right hand to 10%, 20%, and 30% of their maximum voluntary contraction corresponding to low, medium, and high task levels, respectively. Visual cues were provided with Eprime (Version 2.0, Psychology Software Tools, Pittsburgh, PA) using a vertical white bar and a white square bracket that spanned the target force ±1 kgf. The white bar turned green when the force applied to the dynamometer was within the square bracket. The force measures were sent to Eprime real-time using BIOPAC’s network data transfer and a socket connection (BIOPAC, AcqKnowledge Data Acquisition and Analysis Software). Immediately prior to starting the force matching experiment, maximum voluntary contraction was measured by asking the participant on the scanner bed to squeeze the dynamometer as hard as possible for a count of three with verbal encouragement.

For the finger tapping task, participants responded by pressing buttons on an MRI-compatible five button fiber optic response pad (Pyka 5 button handheld, Current Designs, Inc. PA, USA) with their right hand. Visual cues were provided with Eprime by placing a white circle at the distal aspect of the digits of a white hand, indicating when and which button to press. The circles appeared at a rate of 1 Hz, remained visible for 900 ms, and turned green if the correct button was pressed. The experiment had three task levels: a single-digit response with the second digit only, a single-digit response with all digits in a sequential order, and a single-digit response with all digits in a random order, corresponding to low, medium, and high task levels, respectively.

The force matching and finger tapping tasks started with an initial 15 s rest period. Then, ten 15 s trials were performed at each task level with a 15 s rest period after each trial. A total of 30 trials were performed in each experiment. The order of the experiments and the task levels within each experiment were pseudorandomized. **Figure 1** summarizes the visual cues for both force matching and finger tapping tasks.

**Figure 1.**
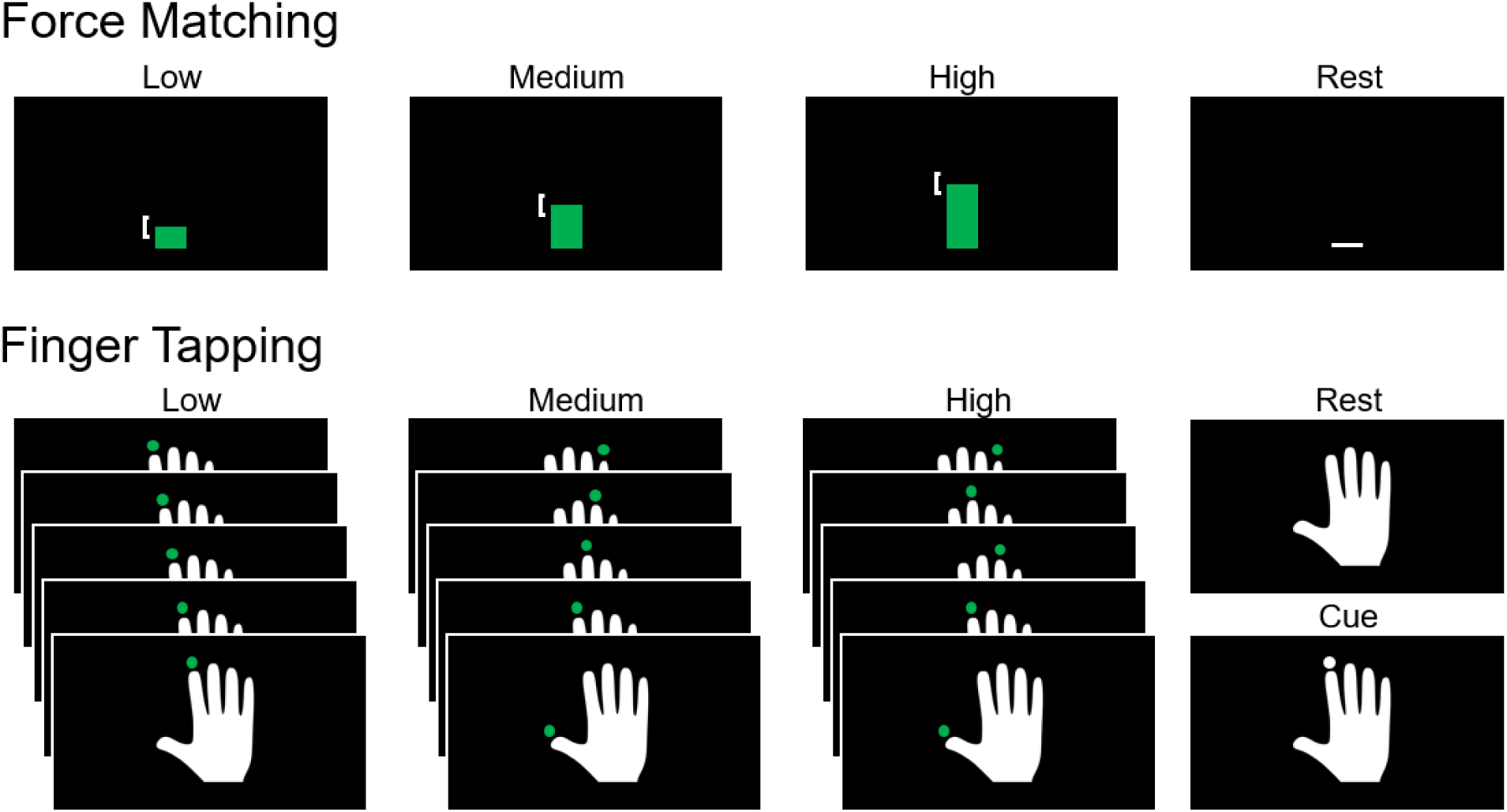
Visual cues for the force matching and finger tapping tasks. For the force matching task, participants gripped an MRI-compatible dynamometer with their right hand to match 10%, 20%, and 30% of maximum voluntary contraction, corresponding to low, medium, and high task levels, respectively. Visual cues were provided with a vertical white bar and a white square bracket that spanned the target force ±1 kgf. The white bar turned green when the force applied to the dynamometer was within the square bracket. For the finger tapping task, participants responded by pressing buttons on an MRI-compatible five button response pad with their right hand. Visual cues were provided by placing a white circle at the distal aspect of the digits of a white hand, indicating when and which button to press. The circles appeared at a rate of 1 Hz, remained visible for 900 ms, and turned green if the correct button was pressed. The experiment had three task levels: a single-digit response with the second digit only, a single-digit response with all digits in a sequential order, and a single-digit response with all digits in a random order, corresponding to low, medium, and high task levels, respectively.

### 2.4. Brain and Spinal Cord Image Acquisition

Brain and spinal cord imaging was performed on a 3T GE SIGNA Premier scanner with a 21-channel neurovascular coil. Participants were briefly trained on the force matching and finger tapping tasks in the scanner room before being placed supine on the scanner bed. Participants were positioned to ensure maximum quality of the spinal cord functional images (Cohen-Adad et al., 2021). The cervical spine was positioned as straight as possible using a cushion below their head to minimize the cervical lordosis. This ensures that axial slices are orthogonal to the spinal cord centerline, thus minimizing partial volume effect (Cohen-Adad et al., 2021). A SatPad^TM^ cervical collar was used to increase the magnetic field homogeneity across the cervical spine and further reduce motion during scanning (Maehara et al., 2014). Physiological data were collected with a pulse oximeter attached to the left index finger and a respiratory belt around the abdomen. Functional imaging was performed using a simultaneous T2*-weighted brain–spinal cord sequence developed by our team (Islam et al., 2019). Briefly, the sequence uses separate field-of-views (FOVs) for the brain (spatial-spectral pulse, 30 slices, full field of view, 3.43 mm ✕ 3.43 mm ✕ 5.00 mm) and cervical spinal cord (2D spatially selective reduced FOV pulse, 15 slices, 1.25 mm ✕ 1.25 mm ✕ 5.00 mm). We aligned the brain FOV axially, while the spinal cord FOV was centered at the C5-C6 intervertebral disc and oriented obliquely such that the slices were perpendicular to the spinal cord. Due to decreasing signal in the inferior cervical spine from increasing distance from the receive coil and closer proximity to the lungs and associated B_0_ inhomogeneity, the investigation of spinal cord activity was limited to the C5 to C7 spinal cord segments. Axial (FOV = 220 mm ✕ 220 mm, matrix size = 128 ✕ 128, ΔTE = 0.5 ms) and sagittal (FOV = 300 mm ✕ 300 mm, matrix size = 256 ✕ 64, ΔTE = 0.5 ms) field maps were collected to calculate the *x* and *y* shim values, while the optimal *z* shim was manually selected by the imaging technologists. Dynamic, x, y, and z shimming was then applied to each brain and spinal cord slice to maximize the signal intensity and minimize distortions. Functional images were acquired with 2.6 s TR (TE = 30 ms, flip angle = 80, bandwidth = 250 kHz, GRAPPA acceleration = 2, phase encoding direction = A/P). A two volume acquisition with opposite phase-encoding direction was collected immediately following the functional runs for distortion correction of the brain functional images. Finally, T1-weighted brain image (3D FSPGR sequence, 1.0 mm ✕ 1.0 mm ✕ 1.0 mm) and a T2-weighted spinal cord image were collected (3D turbo spin-echo, 0.8 mm ✕ 0.8 mm ✕ 0.8 mm) for registration and spatial normalization.

### 2.5. Brain and Spinal Cord Image Processing

Brain and spinal cord functional images for both the force matching and finger tapping tasks were preprocessed and analyzed with separate but analogous pipelines. Preprocessing was performed using commands from Advanced Normalization Tools (ANTS) (version=2.5.1) (Avants et al., 2011), The Oxford Center for fMRI of the Brain’s (FMRIB) Software Library (FSL) (version = 6.0) (Jenkinson et al., 2012) and the Spinal Cord Toolbox (SCT) (version=6.2) (De Leener et al., 2017). All preprocessing and analysis code is available on GitHub (https://github.com/kennethaweberii/Neuromuscular_Signature_R01_Pilot/releases/tag/r20241107).

#### 2.5.1. Brain Preprocessing

The T1-weighted structural image was brain extracted using `antsBrainExtraction` (Avants et al., 2011), and then `fast` was used to segment the brain-extracted image into gray matter (GM), white matter (WM), and cerebrospinal fluid (CSF) (Zhang et al., 2001). The functional images were motion corrected with `mcflirt` before correction for susceptibility induced distortions using `topup`. The WM and CSF masks, generated from the segmentation of the T1-weighted brain image, were initially registered to the functional images using `flirt` (Jenkinson & Smith, 2001), and the first principal components from each were extracted to create volume-wise WM and CSF nuisance regressors with `fslmeants`. Slice-wise cardiac and respiratory noise regressors were also generated using `pnm` (Brooks et al., 2008) for a total of 32 regressors. These regressors were then included in a general linear model using `feat` along with the six motion parameters to remove WM, CSF, physiological, and motion-related noise from the brain functional time series.

The mean brain functional image was co-registered to the T1-weighted structural images using `flirt` with six degrees of freedom, and then the brain-extracted T1-weighted structural image was normalized to the MNI152 template space (2 mm ✕ 2 mm ✕ 2 mm) using non-linear registration with 12 degrees of freedom and a 10 mm warp field in `fnirt` (Andersson et al., 2007.). These transformations were concatenated, and the functional times series was then warped to standard space using `applywarp.` The functional images were then high-pass temporally filtered (100 s cut-off) using `fslmaths` and spatial smoothed with a 5 mm ✕ 5 mm ✕ 5 mm full width half maximum (FWHM) with `susan`. These preprocessing steps generated denoised functional brain images in a standard space to be used as inputs for the first level analyses.

#### 2.5.2. Spinal Cord Preprocessing

Motion correction of the functional spinal cord time series was first performed using `flirt`. A mask around the spinal canal was generated by combining and dilating the automatic segmentations of the CSF and spinal cord output by `sct_propseg` and used to exclude areas outside of the spinal column (De Leener et al., 2014). The functional time series were then aligned with the mid-volume of the functional time series using two-dimensional rigid realignment of each axial slice, the mean image was then calculated, and the motion correction realignment was repeated aligning each volume to the mean image (Weber et al., 2014). At each step of motion correction, the functional time series was visually inspected.

The spinal cord was then segmented from the mean functional image of the motion-corrected time series with `sct_deepseg` (Bédard et al., 2025). A spinal canal mask was generated by combining the spinal cord and CSF segmentations output from `sct_propseg`, and a CSF mask was generated by subtracting the `sct_deepseg` spinal cord segmentation from the spinal canal segmentation. All segmentations were visually assessed using `sct_qc and` manually corrected as necessary.

The spinal cord from the T2-weighted structural image was segmented automatically using the using SCT’s contrast-agnostic segmentation method, `sct_deepseg` (Bédard et al., 2025), and the vertebral levels were automatically labeled using `sct_label_vertebrae` (Ullmann et al., 2014). The spinal cord T2-weighted image was then registered to the PAM50 spinal cord template using `sct_register_to_template` (De Leener et al., 2018). The PAM50 template was registered to the mean spinal cord functional image with `sct_register_multimodal` using the spinal cord segmentations, non-linear slicewise transforms, and the T2-weighted structural image to PAM50 template transformation to initialize the registration. These transformations were concatenated, the PAM50 template was warped to the functional images using `sct_warp_template`, and a WM mask was generated by thresholding and binarizing the PAM50 WM probability atlas (threshold = 0.9).

Slice-wise WM and CSF nuisance noise regressors were generated by extracting the first principal components of the WM and CSF time series, respectively, with `fslmeants`. Slice-wise cardiac and respiratory noise regressors were generated using `pnm` (32 regressors) as well as slice-wise motion regressors (x translation, y translation, and z rotation). These regressors were included in a general linear model using `feat` to remove physiological- and motion-related noise from the spinal cord functional time series. The denoised functional times series was warped to the PAM50 template. In the template space, smoothing was applied with a 2 mm ✕ 2 mm ✕ 5 mm FWHM Gaussian kernel, to preferentially smooth along the superior-inferior axis of the spinal cord. These preprocessing steps generated denoised functional spinal cord images in a standard space to be used as inputs for the subject level analyses. The temporal signal-to-noise ratio (tSNR) was similar for both tasks (**Figure S1**).

#### 2.5.3. Subject Level Analysis

Subject level analyses were performed identically for brain and spinal cord data. The force matching and finger tapping tasks were modeled using trialwise boxcar regressors convolved with a double gamma hemodynamic response function. The trialwise task activation maps from the preprocessed functional time series were then entered into a second level fixed effects model to create average subject level activity maps for each task level. A linear contrast across the task levels was also applied to map where the activity scaled linearly with task level.

#### 2.5.4. Group Level Analysis

The second level activity brain maps were then entered into a mixed effects analysis (FLAME Stage 1) to generate group level activity maps with age and sex included as covariates of no interest. The group level brain activity was defined using a voxelwise threshold of Z score > 3.10 with a cluster significance threshold of p < 0.05 to correct for multiple comparisons. *A priori* a mixed effects analysis was planned for the group level spinal cord analysis. However, significant group level spinal cord activity was not consistently present with a mixed effects analysis, and instead, the reported group level spinal cord activity maps were generated from a fixed effects analysis, and group level spinal cord activity was defined using a voxelwise threshold of Z score > 1.64 with a cluster significance threshold of p < 0.05 to correct for multiple comparisons. The number of active voxels and the average Z score of the active voxels were extracted from each task level at the group level to assess the spatial extent and magnitude of the activity. Both activations (i.e., positive signal change) and deactivations (i.e., negative signal change) were assessed. For the spinal cord, analyses were confined to the region of intersection of the subject level maps, which spanned the C5 to C7 spinal cord segment levels.

### 2.6. Spinal Cord Spatial Analysis

To summarize the localization of the activity, left-right (LR) indices at the group level were calculated by dividing the difference in the number of active voxels between the respective hemicords by the sum (number of active voxels in the entire spinal cord) (Seghier, 2008). A value of +1.00 indicates that all active voxels were in the left hemicord while a value of –1.00 indicates that all active voxels were in the right hemicord. The localization of the activity to the GM and WM was also assessed. As the volume of the WM is more than three times the volume of the GM, the ratio of the percentage of GM activation to the percentage of WM, GM-WM ratio, was calculated to account for the differences in volume. LR indices and GM-WM ratios were assessed at the group level for each task level as well as the linear contrast.

### 2.7. Region of Interest Analysis

To explore local changes in the magnitude of activity across the three task levels, we performed a brain and spinal cord region of interest (ROI) analysis. For the brain ROI’s, spheres (radius = 5 mm) were placed at the supplementary motor area (SMA), dorsal premotor cortex (dPMC), ventral premotor cortex (vPMC), primary motor cortex (M1), and primary somatosensory cortex (S1), regions responsible for human motor movement (Tanji, 2001). The selection of coordinates for the brain ROI’s was initially guided by the meta-analysis of Mayka et al. (Mayka et al., 2006) and refined using the Mindboggle-101 DKT31 cortical atlas (Klein & Tourville, 2012) and Neurosynth (Yarkoni et al., 2011) association test maps (FDR < 0.01 thresholded) for the following terms: hand movements, supplementary motor, premotor, primary somatosensory. The brain coordinates in MNI space are provided in the Supplementary Material (**Table S1).** The spinal cord ROI’s included the left and right gray matter from the PAM50 template (threshold > 0.50) spanning the C5 to C7 spinal cord segment levels. The average Z score within each ROI was extracted for each task level. Two-tailed one-sample t-tests were performed for each task level to identify positive (i.e., activation) or negative (i.e., inactivity) activity within each ROI. Two-tailed paired t-tests were performed between each task level to identify differences in activity between task levels for each ROI. Repeated measures, mixed effects linear models were performed to explore associations between left M1 activity and spinal cord GM activity across the task levels. Statistical significance for these exploratory analyses was set at an α < 0.05 without correction for multiple comparisons. Statistical analyses were performed in R via rpy2 (*rpy2: Python-R Bridge*).

### 2.8. Trialwise Analysis

For each subject and task level, the task error, the number of active voxels, and average Z score of the active voxels were extracted for each trial. For the force matching task, task error was assessed using the mean absolute percent error, which was calculated for each subject and trial for the low, medium, and high task levels. For the finger tapping task, task error was calculated as the percentage of missed taps in each trial for the low, medium, and high task levels. Repeated measures, mixed effects linear models were performed to explore linear changes in activity and task errors across the runs to identify the presence of fatigue or motor learning across the experiments. Each task level was investigated separately. Statistical significance for these exploratory analyses was set at an α < 0.05 without correction for multiple comparisons.

## 3. Results

### 3.1. Hand Function

All participants were right-handed based on the Edinburgh handedness short form (range = +33 – +100 kgf). Average right hand grip strength was 22.1 ± 11.2 kgf (range = 6.3 – 48.8 kgf), and average NIH Toolbox dominant hand grip strength t-score was 45.2 ± 7.9 (range = 28 – 58). The average NIH Toolbox dominant hand dexterity t-score was 58.6 ± 10.5 (range = 37 – 77). The average Quick DASH outcome score was 3.2 ± 5.0 (range = 0 – 22.7), and the average PROMIS Upper Extremity score was 53.9 ± 5.3 (range = 40.6 – 57.1).

### 3.2. Force Matching Task

Brain activity maps show activations and deactivations in cortical and subcortical motor and sensory areas at each task level (mixed effects, p < 0.001, cluster correction threshold p < 0.05) for the force matching task (**Figure 2**). At the group level, the number of active voxels for the activations and deactivations was highest in the high task level. The average Z score of the activations and deactivations, on the other hand, decreased with increasing task level.

**Figure 2.**
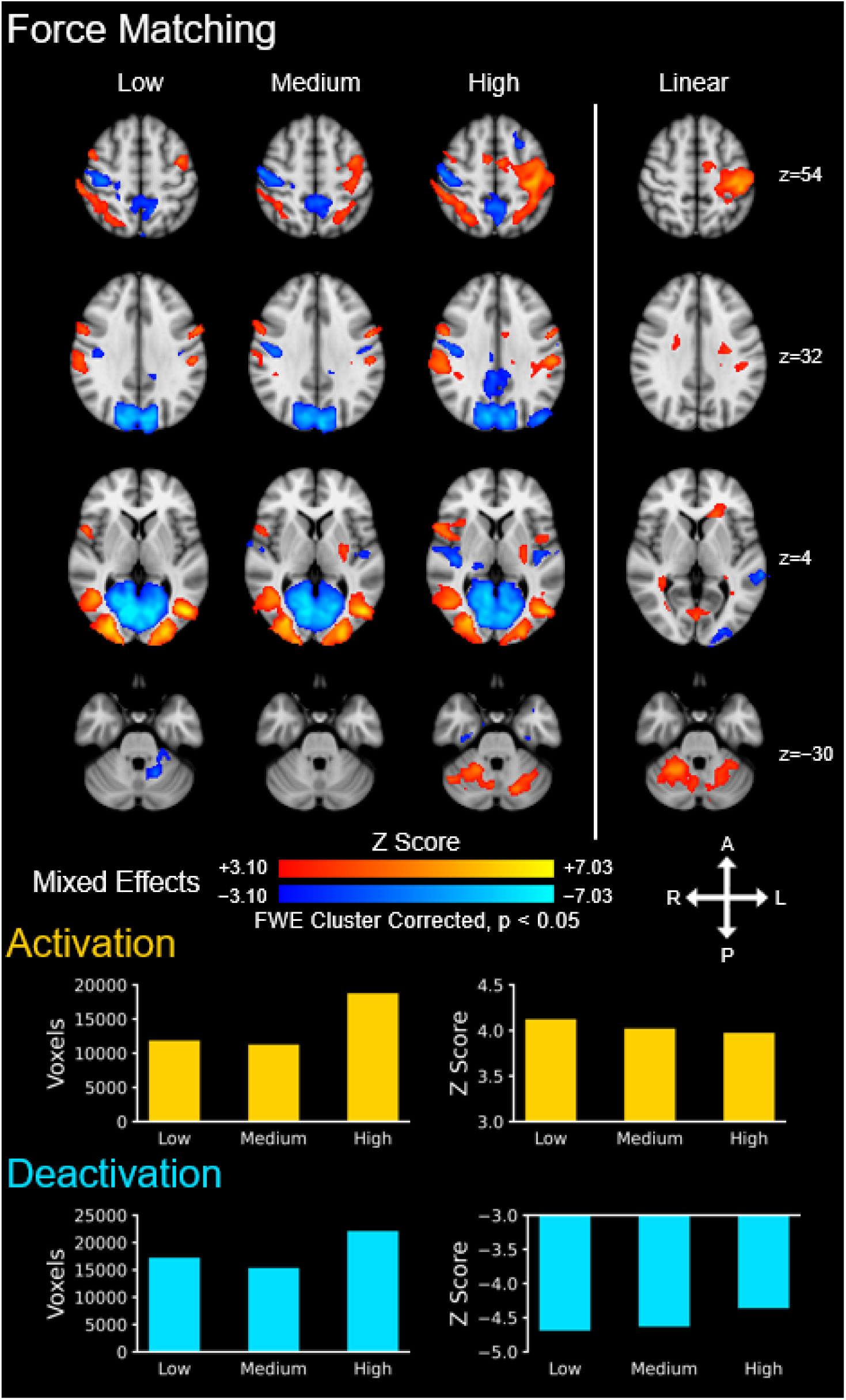
Group level brain activity for the force matching task across the three task levels: low, medium, and high. Activations (i.e., positive signal change) are shown in red–yellow and deactivations are shown in blue–light blue (i.e., negative signal change). A linear contrast across the task levels was applied to map where the signal linearly increases and decreases across the task levels. The number of active voxels and the average Z score of the active voxels are shown to summarize the spatial extent and magnitude of the activity across the three task levels. The activation maps were generated from a mixed effects analysis at the group level and were voxel-wise thresholded at a Z score > 3.09 with a family-wise error (FWE) cluster correction threshold of p < 0.05. The background image is the MNI152 T1-weighted brain template. A = anterior, P = posterior, L = left, R = right.

Spinal cord activity maps for the force matching task show activation in the right ventral and dorsal horns for each task level (i.e., negative LR indices), and deactivation in the left ventral and dorsal horns only at the high task level (**Figure 3)**. Activation in the right ventral and dorsal horns linearly increased with increasing task level while deactivation within the left ventral and dorsal horns linearly increased with increasing task level (fixed effects, p < 0.05, cluster correction threshold p < 0.05). The activations and deactivations were localized more in the gray matter (i.e., positive GM-WM ratios) in the high, medium and linear contrasts. At the group level, the number of active voxels and the average Z score of the active voxels for activation increased across the task levels. See the Supplementary Material for spinal cord activity maps using a mixed effects analysis as well maps with and without cluster correction (**Figure S4**, **Figure S6**, and **Figure S8**).

**Figure 3.**
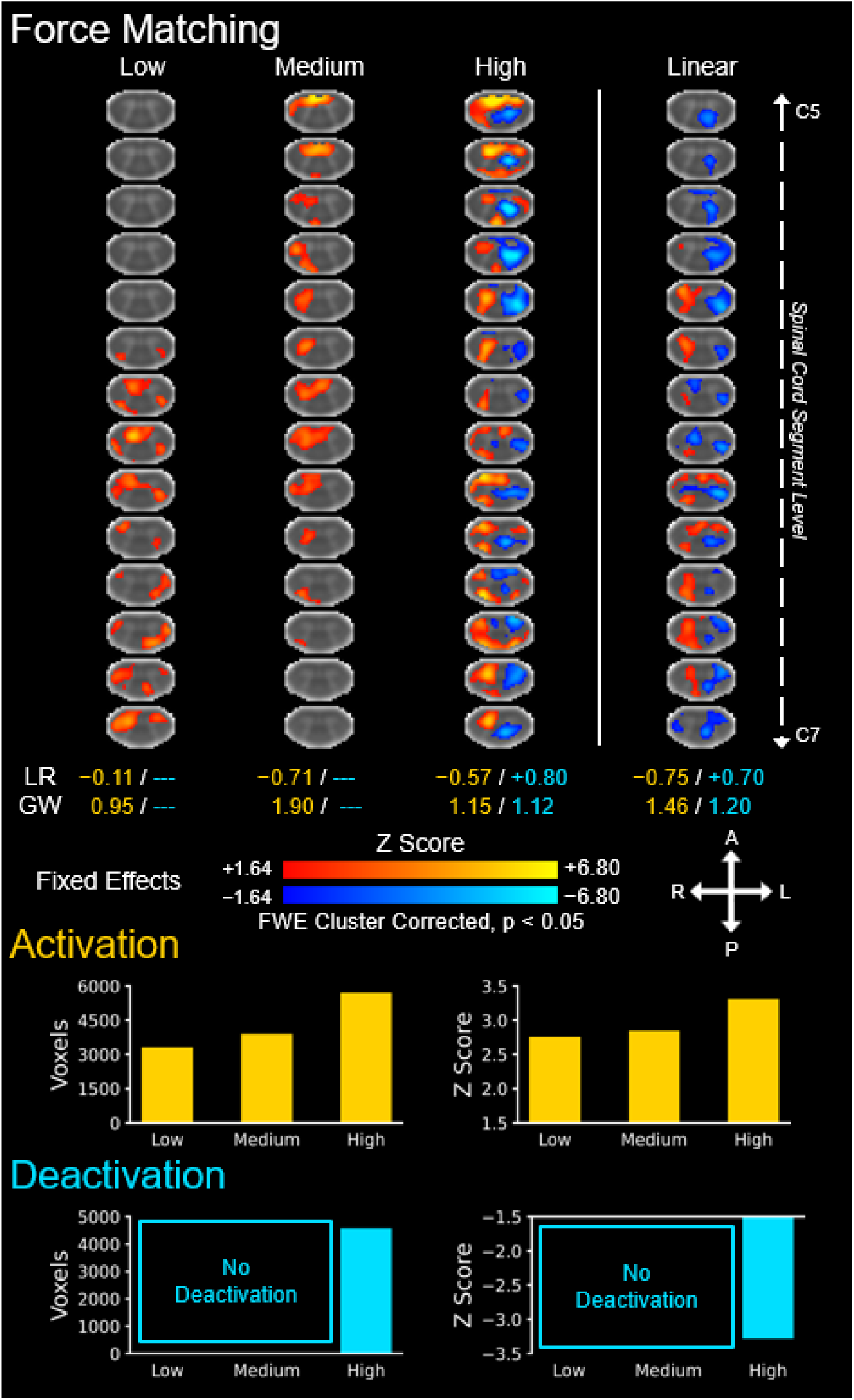
Group level spinal cord activity for the force matching task across the three task levels: low, medium, and high. Activations (i.e., positive signal change) are shown in red–yellow and deactivations are shown in blue–light blue (negative signal change). A linear contrast across the task levels was applied to map where the signal linearly increases and decreases across the task levels. The location of the activations and deactivations was assessed using the left-right (LR) index and gray matter-white matter (GW) ratio (--- = no activity, unable to calculate). The number of active voxels and the average Z score of the active voxels are shown to summarize the spatial extent and magnitude of the activations and deactivations across the three task levels. The number of active voxels and the average Z score of the active voxels are shown to summarize the spatial extent and magnitude of the activity across the three task levels. The activation maps were generated from a fixed effects analysis at the group level and were voxel-wise thresholded at a Z score > 1.64 with a family-wise error (FWE) cluster correction threshold of p < 0.05. The background image is the PAM50 T2*-weighted spinal cord template. Every 5th axial slice from the intersection of the subject level functional images is shown. A = anterior, P = posterior, L = left, R = right.

ROI analysis reveals bilateral activation of the left and right SMA, present across all task levels with the exception of the right SMA during the medium task level (**Figure 4**). SMA activation was highest during the high task level. The left dPMC showed a graded increase in activation across the task levels while the right dPMC was only activated during the high task level. The left and right vPMC were activated at a largely consistent magnitude across the task levels. The left M1 and S1 were activated during the medium and high task levels with a graded increase in activation across the task levels. Interestingly, the right M1 and S1 showed a consistent magnitude of deactivation across the task levels, consistent with contralateral inhibition. The right spinal cord gray matter showed a trend towards activation for the medium and high task levels but the magnitude of activation did not reach statistical significance. On the other hand, deactivation was seen in the left spinal cord gray matter during the high task level. A positive trend between left M1 activity and right spinal cord GM activity was present but did not reach statistical significance (β = 0.217 ± 0.111, t = 1.954, p = 0.054), and a negative association between left M1 activity and left spinal cord GM activity was observed (β = -0.276 ± 0.108, t = -2.553, p = 0.013).

**Figure 4.**
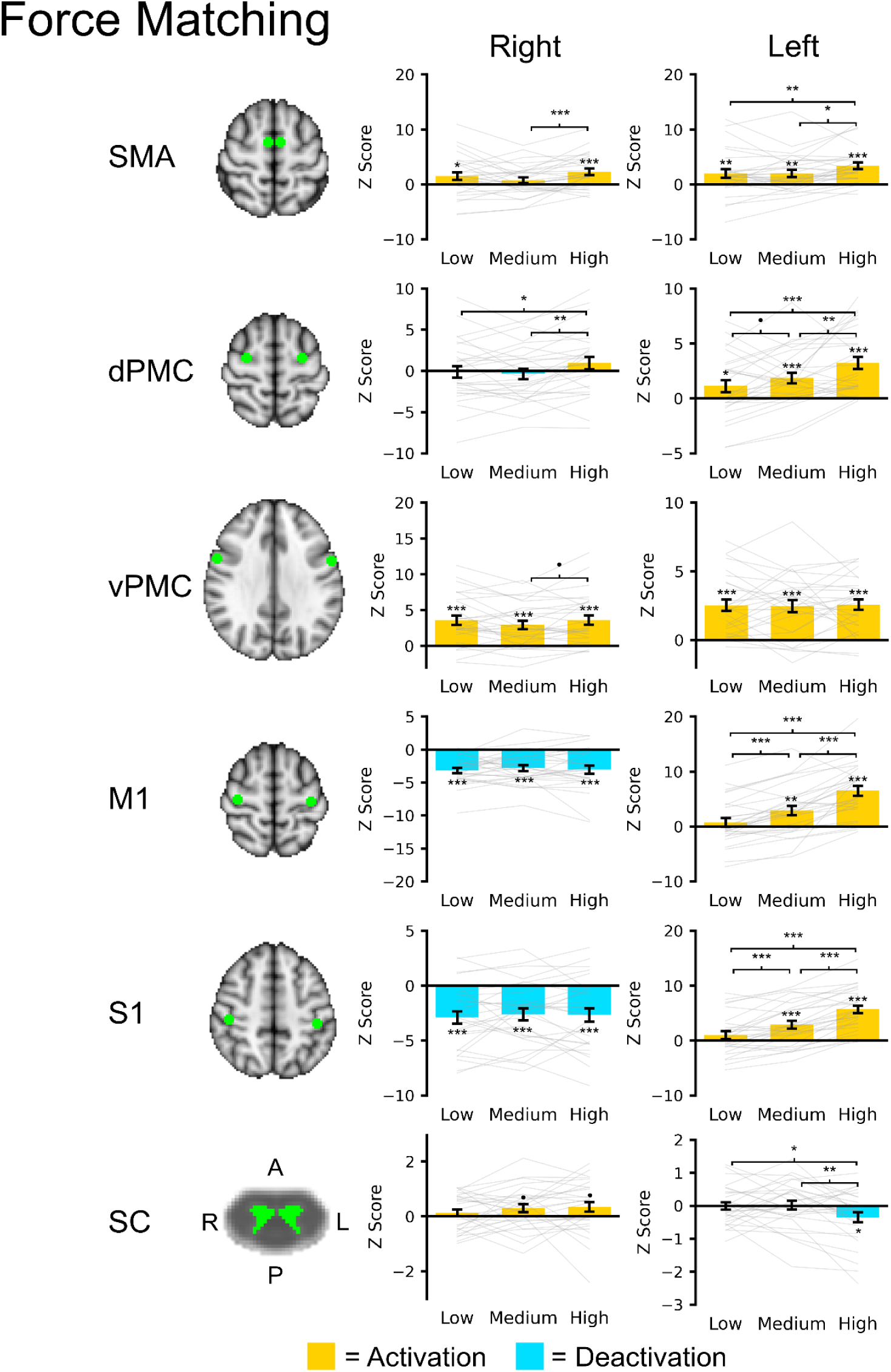
Brain and spinal cord (SC) region of interest (ROI) analysis for the force matching task across the three task levels: low, medium, and high. For the brain ROI’s, spheres (radius = 5 mm) were placed at the supplementary motor area (SMA), dorsal premotor cortex (dPMC), ventral premotor cortex (vPMC), primary motor cortex (M1), and primary somatosensory cortex (S1). The spinal cord ROI’s included the left and right gray matter spanning the C5 to C6 spinal cord segment levels. The brain and spinal cord regions are shown in green overlaid the respective template. The average Z score within each ROI was extracted. Bar plots show the average activation (yellow) or deactivation (blue) within each region across the subjects. Gray lines show subject level activity level across the task levels. Error bars ± 1 SE. •p < 0.10, *p < 0.05, **p < 0.01, and ***p < 0.001.

### 3.3. Finger Tapping Task

Brain activity maps for the finger tapping task are presented in **Figure 5**. Activations and deactivations were seen in cortical and subcortical motor and sensory areas at each task level (mixed effects, p < 0.001, cluster correction threshold p < 0.05). At the group level, the number of active voxels and the average Z score of the activations increased across the task levels. The number of active voxels for the deactivations was lowest for the high task level, and the average Z score of the deactivations decreased across the task levels.

**Figure 5.**
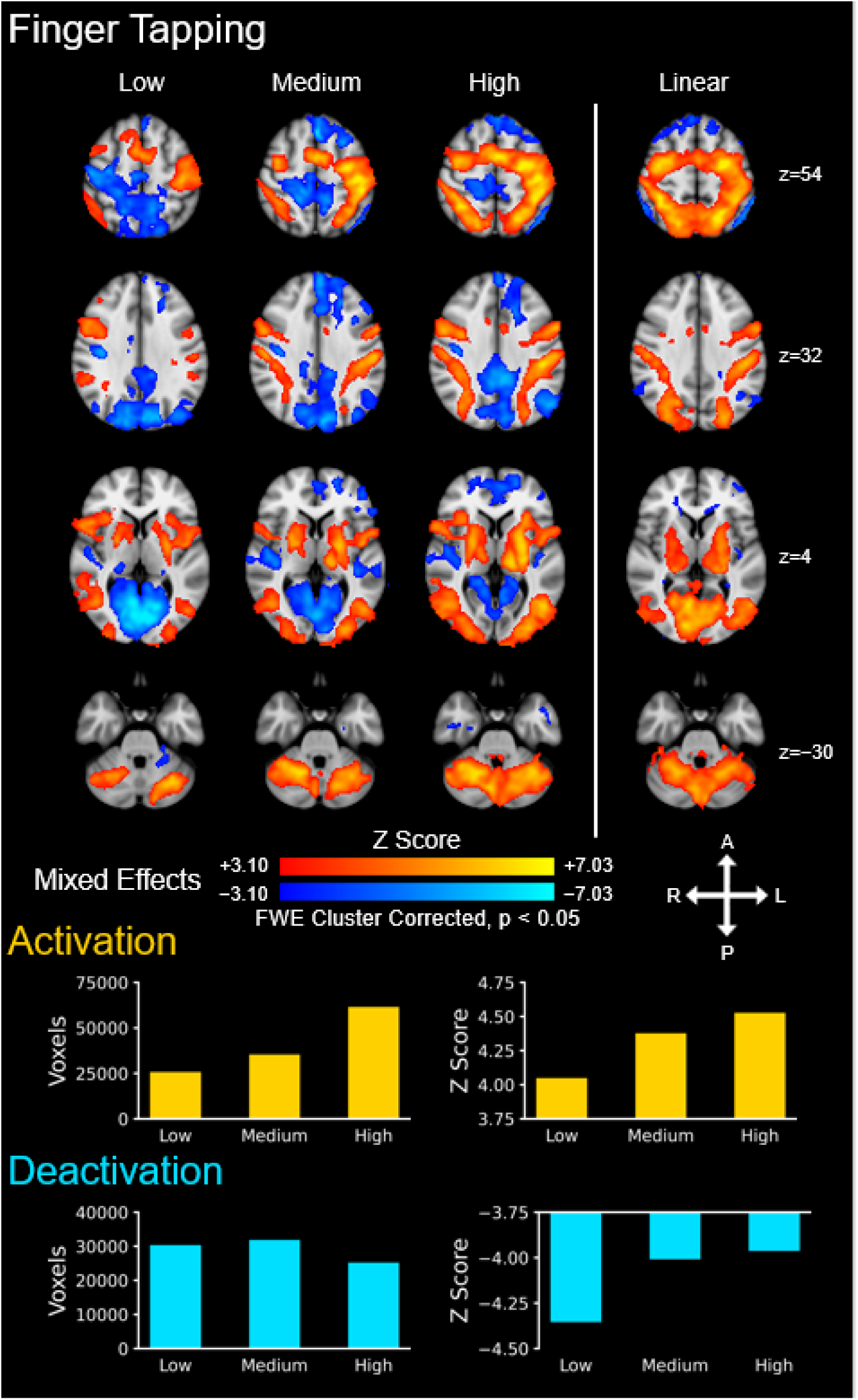
Group level brain activity for the finger tapping task across the three task levels: low, medium, and high. Activations (i.e., positive signal change) are shown in red–yellow and deactivations are shown in blue–light blue (i.e., negative signal change). A linear contrast across the task levels was applied to map where the signal linearly increases and decreases across the task levels. The number of active voxels and the average Z score of the active voxels are shown to summarize the spatial extent and magnitude of the activity across the three task levels. The activation maps were generated from a mixed effects analysis at the group level and were voxel-wise thresholded at a Z score > 3.09 with a family-wise error (FWE) cluster correction threshold of p < 0.05. The background image is the MNI152 T1-weighted brain template. A = anterior, P = posterior, L = left, R = right.

Spinal cord activity maps for the finger tapping task are presented in **Figure 6**. Activation was seen in the right ventral and dorsal horns for each task level (i.e., negative LR indices). No deactivation was present. Activation in the right ventral and dorsal horns linearly increased with increasing task level (fixed effects, p < 0.05, cluster correction threshold p < 0.05). The activations were localized more to the gray matter (i.e., positive GM-WM ratios) for all task levels and the linear contrast. At the group level, the number of active voxels and the average Z score of the active voxels increased across the task levels. See the Supplementary Material for spinal cord activity maps using a mixed effects analysis as well maps with and without cluster correction (**Figure S5**, **Figure S7**, and **Figure S9**).

**Figure 6.**
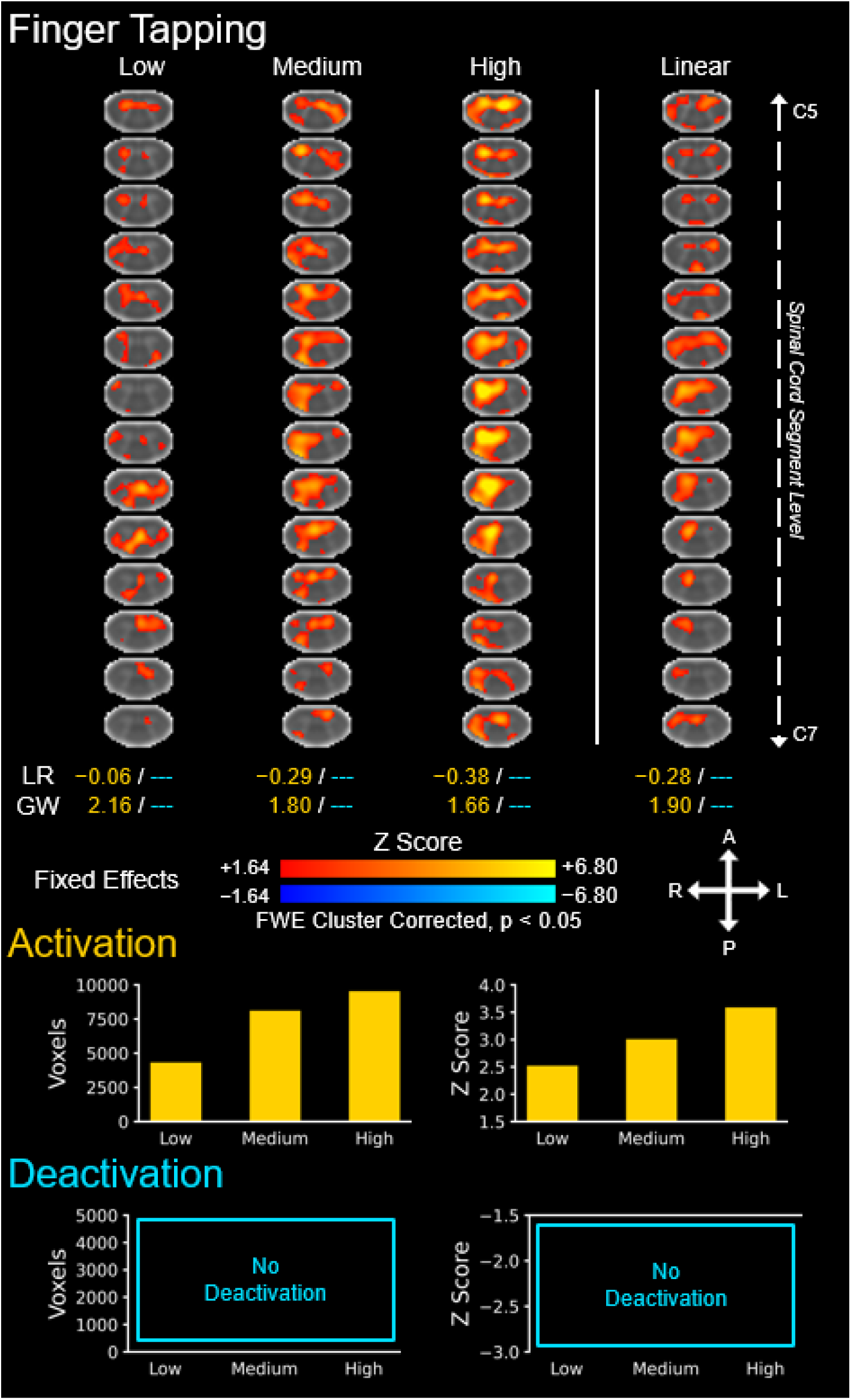
Group level spinal cord activity for the finger tapping task across the three task levels: low, medium, and high. Activations (i.e., positive signal change) are shown in red–yellow and deactivations are shown in blue–light blue (negative signal change). A linear contrast across the task levels was applied to map where the signal linearly increases and decreases across the task levels. The location of the activations and deactivations was assessed using the left-right (LR) index and gray matter-white matter (GW) ratio (--- = no activity, unable to calculate). The number of active voxels and the average Z score of the active voxels are shown to summarize the spatial extent and magnitude of the activations and deactivations across the three task levels. The number of active voxels and the average Z score of the active voxels are shown to summarize the spatial extent and magnitude of the activity across the three task levels. The activation maps were generated from a fixed effects analysis at the group level and were voxel-wise thresholded at a Z score > 1.64 with a family-wise error (FWE) cluster correction threshold of p < 0.05. The background image is the PAM50 T2*-weighted spinal cord template. Every 5th axial slice from the intersection of the subject level functional images is shown. A = anterior, P = posterior, L = left, R = right.

Results from the ROI analysis are presented in **Figure 7**. The left and right SMA showed a graded increase in activation across the task levels. The right dPMC deactivated during the low task level while the left and right dPMC showed a graded increase in activation across the medium and high task levels. The left and right vPMC also showed a graded increase in activation across task levels. The left M1 and S1 were activated during each of the task levels with higher activation during the medium and high task levels compared to the low task levels but no statistical difference in activation between the medium and high task levels. The right M1 and S1 showed deactivation for the low task and no activation for the medium task level. The right S1 but not the right M1 was activated during the high task level. The right spinal cord gray matter showed a trend towards activation for the low task level but the magnitude did not reach statistical significance. For the medium and high task levels, the right spinal cord gray matter was activated with no statistical difference in activation between the medium and high task levels. The left spinal cord gray matter showed a trend towards activation for the medium and high task levels but the magnitude did not reach statistical significance. A positive association between left M1 activity and right spinal cord GM activity was present (β = 0.228 ± 0.108, t = 2.112, p = 0.038) while no significant association between left M1 activity and left spinal cord GM activity was observed (β = -0.037 ± 0.107, t = -0.345, p = 0.731).

**Figure 7.**
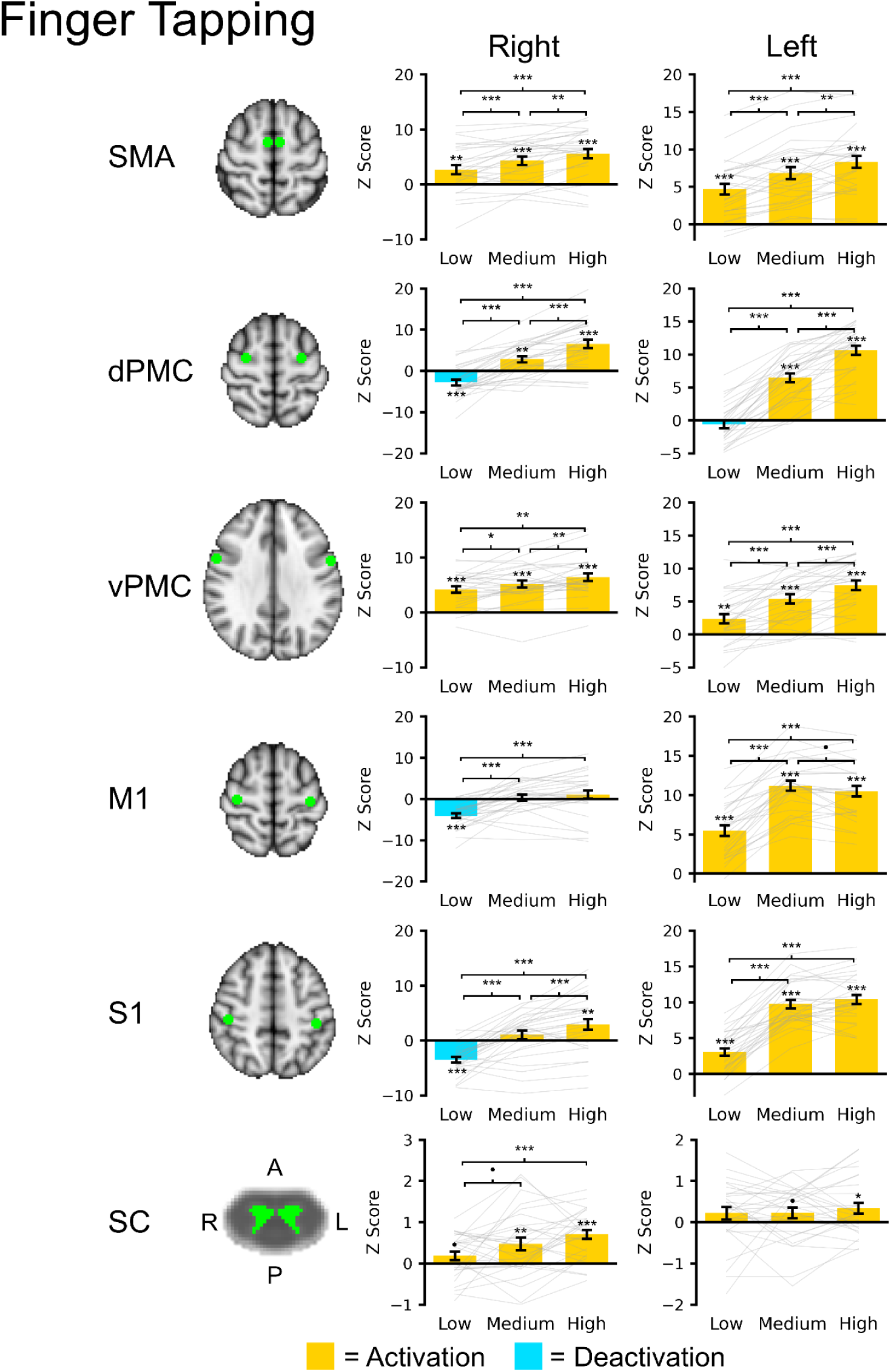
Brain and spinal cord (SC) region of interest (ROI) analysis for the finger tapping task across the three task levels: low, medium, and high. For the brain ROI’s, spheres (radius = 5 mm) were placed at the supplementary motor area (SMA), dorsal premotor cortex (dPMC), ventral premotor cortex (vPMC), primary motor cortex (M1), and primary somatosensory cortex (S1). The spinal cord ROI’s included the left and right gray matter spanning the C5 to C6 spinal cord segment levels. The brain and spinal cord regions are shown in green overlaid the respective template. The average Z score within each ROI was extracted. Bar plots show the average activation (yellow) or deactivation (blue) within each region across the subjects. Gray lines show subject level activity level across the task levels. Error bars ± 1 SE. •p < 0.10, *p < 0.05, **p < 0.01, and ***p < 0.001.

### 3.4. Trialwise Analysis

No consistent linear changes in task error or activity, suggesting the presence of fatigue or motor learning, were identified across the experiments. For the force matching medium task level, task error linearly increased (p = 0.028), and the average Z score of the active voxels in the spinal cord linearly decreased (p = 0.042). No changes in brain activity were identified in the medium task level, and no changes in task error or activity were identified in the low and high task levels (**Figure S2**). For the finger tapping task, no changes in task error were present across the task levels. For the finger tapping medium task level, the number of active brain voxels linearly decreased (p = 0.029) and the average Z score of the active spinal cord voxels linearly decreased (p = 0.003). No changes in brain or spinal cord activity were identified in the low and high task levels (**Figure S3**).

## 4. Discussion

Our study provides novel insights into the integrated neural mechanisms underlying hand function by leveraging simultaneous brain-spinal cord fMRI. Our findings demonstrate that task difficulty modulates both cortical and spinal activation, revealing a dynamic interplay between these regions. Specifically, graded activation in M1 and S1, alongside corresponding responses in spinal gray matter, suggests that motor control involves coordinated top-down modulation rather than isolated cortical command. Additionally, observed spinal deactivations may reflect inhibitory mechanisms essential for fine-tuned motor output. These findings have important mechanistic and clinical implications.

These findings align with previous work demonstrating force-dependent M1 activation (Braaß et al., 2023; Cramer et al., 2002; Dai et al., 2001). The ROI analysis complemented the group level maps showing increasing activation of the left M1 and S1 with increasing force generation. M1 is the main brain region for descending motor commands, which makes monosynaptic connections to motor neurons in the spinal cord ventral horn, via the corticospinal tract, leading to muscle contraction. Motor units (a motor neuron and associated muscle fibers) are recruited in a graded fashion based on size and force production where smaller, lower force generating motor units are recruited first followed by larger, higher force generating motor units (i.e., Henneman’s size principle (Henneman, 1957)). The left M1 activity scaled with force to produce the descending drive needed to recruit the appropriate motor units and achieve the target force levels. Interestingly, we revealed that S1 also scaled with force. In addition to receiving ascending sensory information, S1 sends descending projects to the spinal cord to modulate motor activity and sensory processing based on sensory feedback (Karadimas et al., 2020; Ueno et al., 2018). The graded S1 activity represents the increasing tactile and proprioceptive aspects of the tasks as well as descending modulation. For the finger tapping task, the group level brain activity showed graded activation of the left (i.e., contralateral) M1 and S1, which linearly increased across the task levels. The ROI analysis complements the group level maps but shows a clear step in the magnitude of left M1 and S1 activation from the low task level to the medium and high task levels with the magnitude of activation being largely consistent between the medium and high task levels. The increase in M1 and S1 activity between the low task level, second digit only, and medium and high task levels, all digit tapping (sequential or random order), may represent spatial summation within M1 and S1, as all digit tapping requires activation of more motor units to coordinate the movement. Similarly, all digit tapping spatially activates more afferents than single digit tapping, leading to greater S1 activation. A novel interpretation of this finding is that single-digit tapping of the second digit could be driven more by spinal central pattern generators, neural circuits that produce rhythmic motor activity without descending input (Steuer & Guertin, 2019; Zehr et al., 2004). If so, less descending drive would be needed for the second digit tapping than all digit tapping, since the rhythmic would be spinally mediated, and only descending commands to start and stop the tapping would be needed. The SMA, dPMC, and vPMC are higher level motor areas involved in the planning and organization of motor activity (Wong et al., 2015). These regions receive multisensory information and develop a motor plan that is delivered to M1, which in turn activates the specific muscles. The group level brain activity and ROI analysis showed graded activation of the SMA in both the force matching and finger tapping tasks with more pronounced bilateral activation in the finger tapping task. Graded SMA activation has been reported in similar force matching tasks, highlighting its potential role force generation (Galléa et al., 2008). The SMA is involved in planning complex movements and sequence processing including the order and timing of movements, explaining the graded activation in the finger tapping task, where the SMA was highest for tapping in a random order, which requires the most attention and planning (Cona & Semenza, 2017; Nachev et al., 2008). DPMC directs visually-guided behavior and plans movements in response to visual cues while the vPMC is involved in the perception of space, object manipulation, and transforming positions in space to upper limb movements (Bonnard et al., 2007; Davare et al., 2006; Hoshi & Tanji, 2007). During the force matching task, participants were instructed to keep the force bar within ±1 kgf of the target force. Therefore, the target range at the high task level was smaller relative to the target force than at the low task level. The higher task levels required more visual feedback and attention to match the target force level, which could explain the graded left dPMC activity. The dPMC and vPMC showed increasing bilateral activation across the task levels in the finger tapping task, supporting their roles in planning movements to visual cues and spatial perception, respectively. The patterns of SMA, dPMC, and vPMC activity were largely similar bilaterally with the exception of the right dPMC, which was only activated in the high task level of the force matching task. While only a unilateral task was performed here, the bilateral activation may be due to their roles in planning and coordinating complex movements involving both sides of the body.

The right ventral horn of the spinal cord contains the motor neurons that innervate muscles controlling the right hand. For the force matching task, the group level spinal cord activation maps show right ventral and dorsal spinal cord GM activation with evidence of graded activity across the task levels as expected (Braaß et al., 2023; Madi et al., 2001). Results from the ROI analysis, which averaged activity across the C5 to C7 spinal cord segments, are less clear but provide evidence of lateralized right-sided activity with force generation, as expected (Hemmerling et al., 2023; Weber et al., 2016). Averaging across all spinal cord segments could hide activation if the motor activity was not localized to a single spinal cord segment and may explain the weaker correspondence between the group level spinal cord maps and ROI analysis. For the finger tapping task, the group level activation maps and ROI analysis also show graded right ventral and dorsal spinal cord GM activation. In comparison, the spinal cord activation in the finger tapping task appears to be greater than the force matching tasks, which was not expected and in contrast with previous findings (Maieron et al., 2007). A single button press required < 5 kgf, which is less than the low task level target force of the force matching task for any participant, and because motor units are recruited based on their size, we expected greater spinal cord activity for the force matching task. The greater activation may be from the dynamic nature of the task, leading to less adaptation and greater spatial summation of motor and sensory activity. The greater activation could be due to the complexity of the descending commands for increasing task difficulty, leading to a greater spinal cord activation. Left M1 activity was significantly positively associated with the right spinal cord GM activity for the finger tapping task, consistently with known monosynaptic connections between contralateral motor neurons and ipsilateral spinal cord GM activity (Bortoff & Strick, 1993; Morecraft et al., 2023; Natali et al., 2024). Based on the spatial analysis, the activity was more localized to the right spinal cord and the spinal cord GM. In consistency with previous studies, left spinal cord activation was also present, which may indicate interneuronal processing for the coordination or inhibition of bilateral movements (Maieron et al., 2007; Ng et al., 2008; Stroman & Ryner, 2001; Vahdat et al., 2015; Xie et al., 2009). With fMRI, we are limited to an indirect measure of neural activity based on the blood hemodynamics and the blood oxygenation dependent contrast.

Being able to inhibit unwanted movements is necessary for normal function, so we also interrogated deactivations within the brain and spinal cord (Ferbert et al., 1992; Hanajima et al., 2001). Consistently, the group level activation maps and ROI analyses demonstrated deactivation of the right M1 and S1, which is evidence of interhemispheric inhibition to restrict motor output during the unilateral motor tasks. The deactivation of the left spinal cord ventral and dorsal horns (i.e., contralateral motor and sensory spinal cord regions), seen at the high task level and linear contrast, suggests that the inhibition of motor areas may extend to the spinal cord. In the ROI analysis, the left spinal cord GM was deactivated in the high task level. Further, the magnitude of left M1 activity was negatively associated with the left spinal cord GM activity across the task levels, providing additional indirect evidence of contralateral inhibition. Whether this inhibition results from direct descending modulation from the brain or interneuronal inhibition in the cord remains to be interrogated. Corticospinal projections are complex and synapse on not only motoneurons but also sensory neurons and interneurons, and these projections can be unilateral and bilateral (Ueno et al., 2018). For the finger tapping task, no contralateral deactivation passed cluster correction, but based on the uncorrected maps (**Figure S7**), the deactivation, when present, was more contralateral. The phenomenon of contralateral spinal cord inhibition (i.e., interhemicord inhibition) requires further replication and interrogation.

Our study has several limitations. Both the extrinsic and intrinsic hand muscles contributed to the force matching and finger tapping tasks, and both tasks also likely required coactivation of the wrist flexors and extensors to increase wrist stiffness and stabilize the wrist. Therefore, the spinal cord activation is expected to span the C6 to T1 spinal cord segments (Dalley & Agur, 2023; Hemmerling et al., 2023). Notably, the C8 spinal segment is associated with finger flexion and extension, which may not be fully captured within the C5–C7 coverage. While we may have captured the region of peak motor activity, we were not able to fully interrogate the entire cervical enlargement. We have recently developed a 56-channel research-grade head-neck coil, which can improve imaging in the inferior cervical and upper thoracic spinal cord segments. We are unable to determine whether the increased spinal cord signals are due to increased excitatory or inhibitory activity, descending drive, motor neuron activity, sensory feedback, or spinal interneuronal processing. Combining fMRI with electromyography, peripheral nerve stimulation, and transcranial magnetic stimulation could help disentangle the specific neural mechanisms driving spinal cord single change. For the force matching task, previous studies have used higher levels of force. After pilot testing, we chose a lower level of force (≤ 30% of maximum voluntary contraction) to limit fatigue during the 15-minute experiment, and post-hoc analyses did not demonstrate consistent evidence of fatigue. The low level of spinal cord activation seen in the low force matching task level, especially in the ROI analysis, suggests that the activity in the low task level was just above the noise threshold, and starting at slightly higher force level could have improved the detection of low level force activity. Here we modeled the force matching and finger tapping tasks using the ideal task design. Using the measured force, as performed by Hemmerling et al. (2023), or the timing of the button presses as explanatory variables in the first-level analyses could improve the accuracy of the modeling and the detection of brain and spinal cord activity. Additionally, future studies could use electromyography to capture the actual timing and magnitude of the muscle activity, which could be used to characterize the contributions of different muscle groups to the force matching and finger tapping tasks (Khatibi et al., 2022). Bilateral electromyography could also be used to identify or rule out the presence of mirror movements, involuntary movements in the contralateral limb during a unilateral task. Next, we used the PAM50 template in the Spinal Cord Toolbox, and intervertebral disc levels for spatial normalization, which is not ideal as the location of the spinal cord segments (i.e., where the spinal rootlets enter or exit the spinal cord) in relation to the vertebral bodies varies across participants (Cadotte et al., 2015). Matching the participants on the spinal cord neuroanatomy instead of spine anatomy would likely lead to more robust group level analyses. Only the high task level for the finger tapping task passed cluster correction with a mixed effects analysis, and the use of a fixed effects analysis for the spinal cord activity maps is a limitation and reduces the generalizability of the group level activation map findings (**Figure S5**). We are currently developing techniques for automated identification of the spinal rootlets and spinal cord segment-based spatial normalization methods (Valošek et al., 2024), which should improve the correspondence in spinal neuroanatomy across participants and group level inferences.

We believe our study has several notable strengths. Simultaneous brain-spinal cord fMRI and the inclusion of graded force matching and finger tapping tasks allowed for detailed characterization of motor and sensory activity across the central nervous system under varying demands. The findings showing interhemispheric inhibition and spinal cord deactivation during unilateral tasks, offer novel mechanistic insights into hand motor control and its modulation.

Deactivation of the left spinal cord gray matter during the force matching task provides the first evidence that the inhibition of motor areas during a unilateral motor task extends to the spinal cord. Whether this inhibition results from direct descending modulation from the brain or interneuronal inhibition in the spinal cord remains to be interrogated.

In summary, our study advances understanding of brain-spinal cord interactions in motor function by demonstrating graded neural activation and inhibition across the neuraxis. By integrating brain and spinal cord imaging, we provide a comprehensive model of sensorimotor control with direct translational relevance. These findings pave the way for future research on targeted neurorehabilitation approaches aimed at optimizing motor function in individuals with neurological impairments.

## 5. Acknowledgements

Computing for this project was performed on the Sherlock high-performance computing cluster. We would like to thank Stanford University and the Stanford Research Computing Center for providing computational resources and support that contributed to these findings.

## 6. Funding

This study was supported by grants from the National Institute of Neurological Disorders and Stroke (grant numbers K23NS104211, L30NS108301, R01NS128478, and K24NS126781). The content is solely the responsibility of the authors and does not necessarily represent the official views of the National Institutes of Health. Andrew C. Smith was also supported by the Boettcher Foundation’s Webb-Waring Biomedical Research Program and by the Eunice Kennedy Shriver National Institute of Child Health and Human Development (K01HD106928). Valeria Oliva was also supported by the Italian National Institute of Health (Starting Grant for Young Researchers CUP I85E23001090005). Sandrine Bédard is supported by NSERC (the Canada Graduate Scholarships — Doctoral program). Dario Pfyffer is supported by a Swiss National Science Foundation Postdoc.Mobility Fellowship grant (P500PM_214211) and the Redlich Pain Research Endowment.

## 7. Ethics

The study was approved by Stanford University’s Institutional Review Board.

## 8. Data/Code Availability

The code used to analyze the present dataset is available on Github: https://github.com/kennethaweberii/Neuromuscular_Signature_R01_Pilot/releases/tag/r20241107 The data will be made available on openneuro.org.

## 9. Author Contributions

Conceptualization: KW; Methodology, Software and Formal analysis: VO, SB, MK, DP, BC, SA, GG, SM, CL, KW; Investigation: VO, SB, MK, DP, BC, SA, NB, CL, KW; Writing: VO, SB, MK, DP, AC, ST, SH, JR, ZS, AS, GG, SM, CL, KW; Supervision: KW, SM; Funding acquisition: KW

## 10. Competing Interests

The authors declare no competing interests.

## 12. Supplementary Material

**Table S1.** MNI coordinates (x, y, z) of brain regions used in the region of interest analysis.

|  | Right (x, y, z) | Left (x, y, z) |
| --- | --- | --- |
| SMA | 6, -4, 56 | -6, -4, 56 |
| vPMC | 28, -8, 58 | -26, -8, 58 |
| VPM | 56, 6, 32 | -56, 4, 32 |
| M1 | 36, -20, 58 | -36, -22, 58 |
| S1 | 44, -26, 50 | -42, -30, 50 |

**Figure S1.**
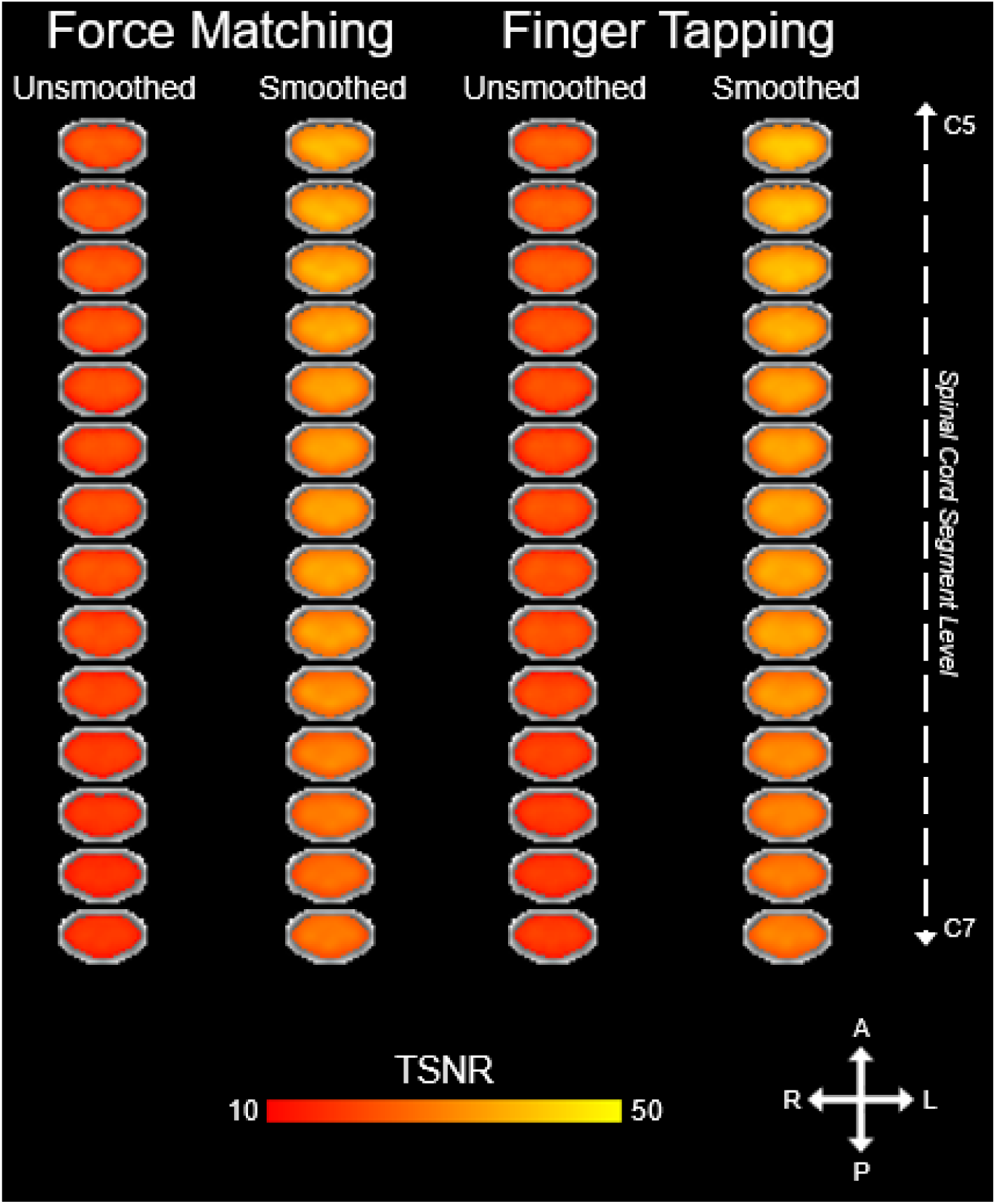
Group level average spinal cord temporal signal-to-noise ratio (TSNR) maps for the force matching and finger tapping tasks with and without spatial smoothing.

**Figure S2.**
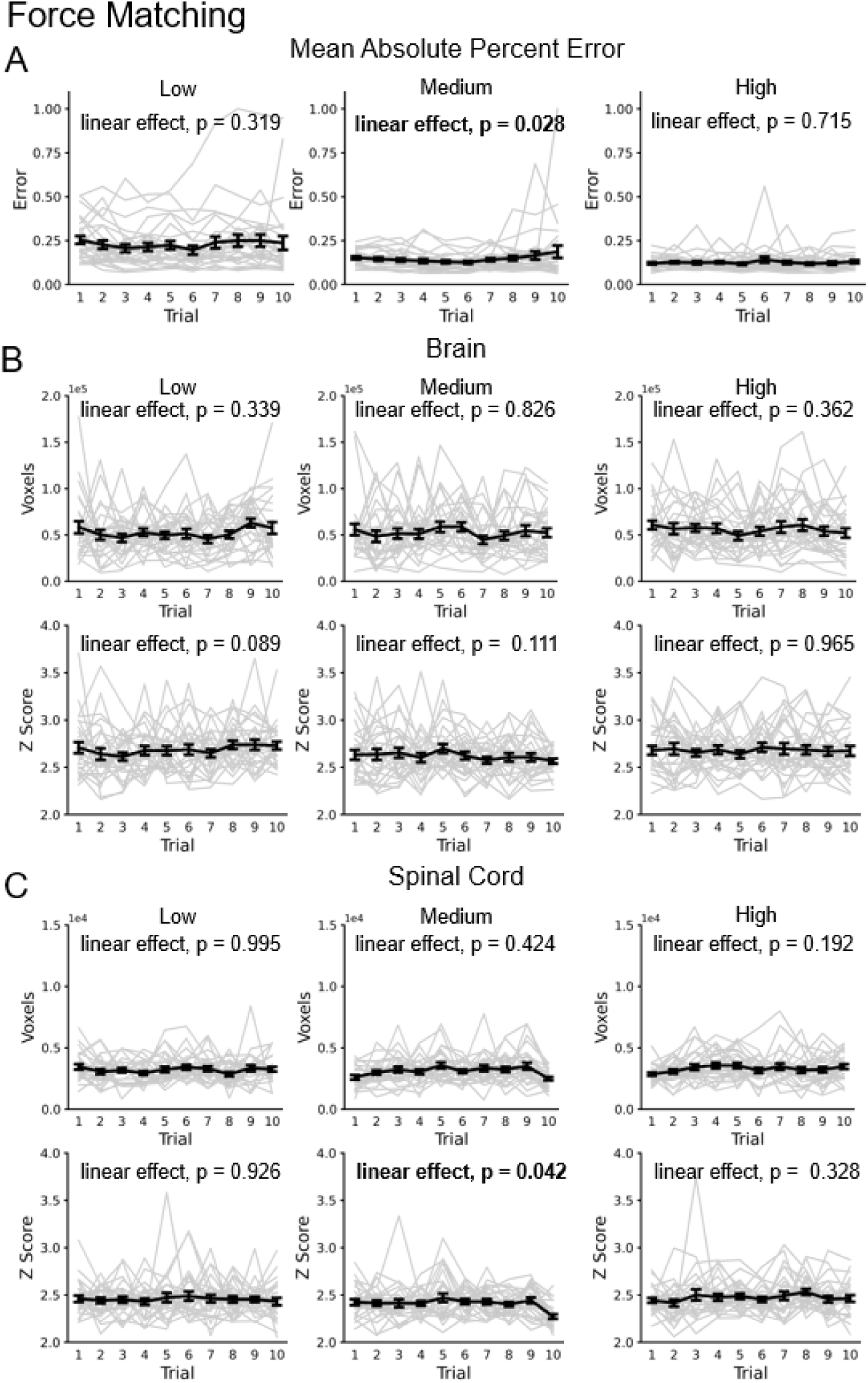
Trialwise error and brain and spinal cord activation plots (number of active voxels and average Z score of the active voxels) from the force matching task for each trial and task level: low, medium, and high. The average values (± one standard error) across the participants are shown in black. While slight linear increases in the mean absolute error for the medium task level and the number of active spinal cord voxels for the high task level were observed, no consistent linear increases or decreases were present across the experiment, indicating that task performance and activity were largely stationary over the course of the force matching task and no strong evidence of fatigue and motor learning was observed.

**Figure S3.**
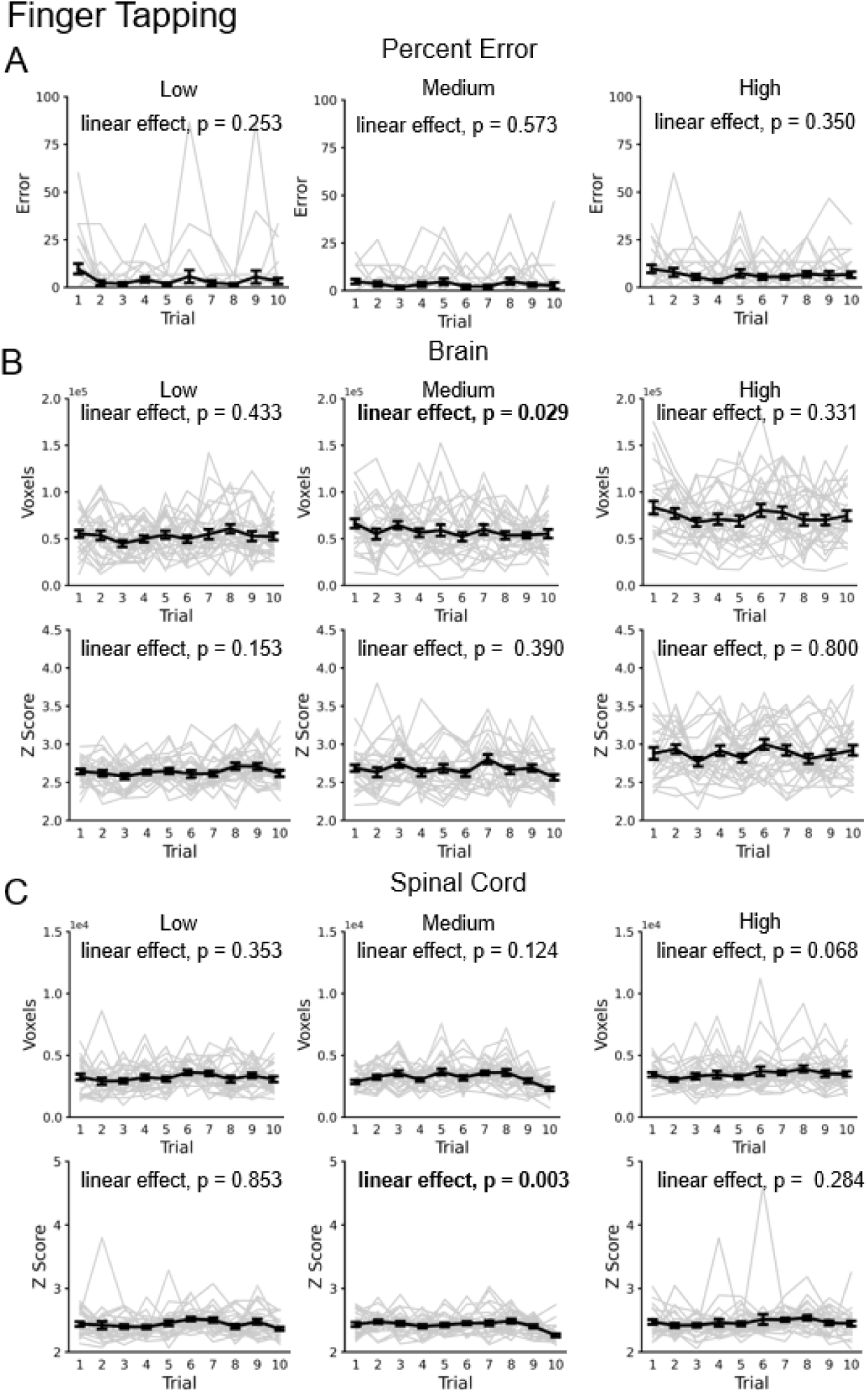
Trialwise error and brain and spinal cord activation plots (number of active voxels and average Z score of the active voxels) from the finger tapping task for each trial and task level: low, medium, and high. The average values (± one standard error) across the participants are shown in black. While slight linear decreases in the number of active brain voxels and the average Z score of the active spinal cord voxels in the medium task level were observed, no consistent linear increases or decreases were present across the experiment, indicating that task performance and activity were largely stationary over the course of the force matching task and no strong evidence of fatigue and motor learning was observed.

**Figure S4.**
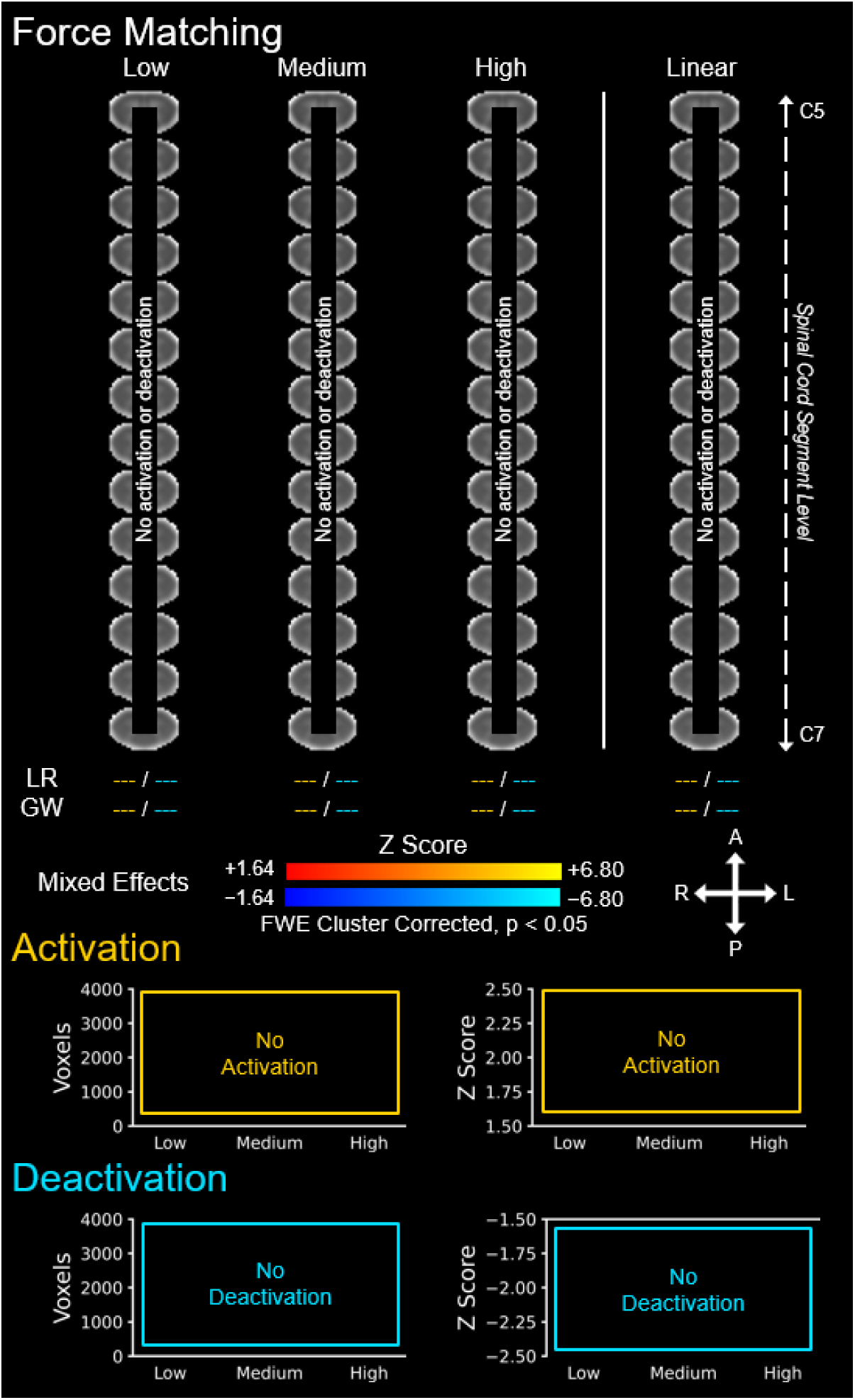
Group level spinal cord activity for the force matching task across the three task levels: low, medium, and high. Activations (i.e., positive signal change) are shown in red–yellow and deactivations are shown in blue–light blue (negative signal change). A linear contrast across the task levels was applied to map where the signal linearly increases and decreases across the task levels. The location of the activations and deactivations was assessed using the left-right (LR) index and gray matter-white matter (GW) ratio (--- = no activity, unable to calculate). The number of active voxels and the average Z score of the active voxels are shown to summarize the spatial extent and magnitude of the activations and deactivations across the three task levels. The number of active voxels and the average Z score of the active voxels are shown to summarize the spatial extent and magnitude of the activity across the three task levels. The activation maps were generated from a mixed effects analysis at the group level and were voxel-wise thresholded at a Z score > 1.64 with a family-wise error (FWE) cluster correction threshold of p < 0.05. The background image is the PAM50 T2*-weighted spinal cord template. Every 5th axial slice from the intersection of the subject level functional images is shown. A = anterior, P = posterior, L = left, R = right.

**Figure S5.**
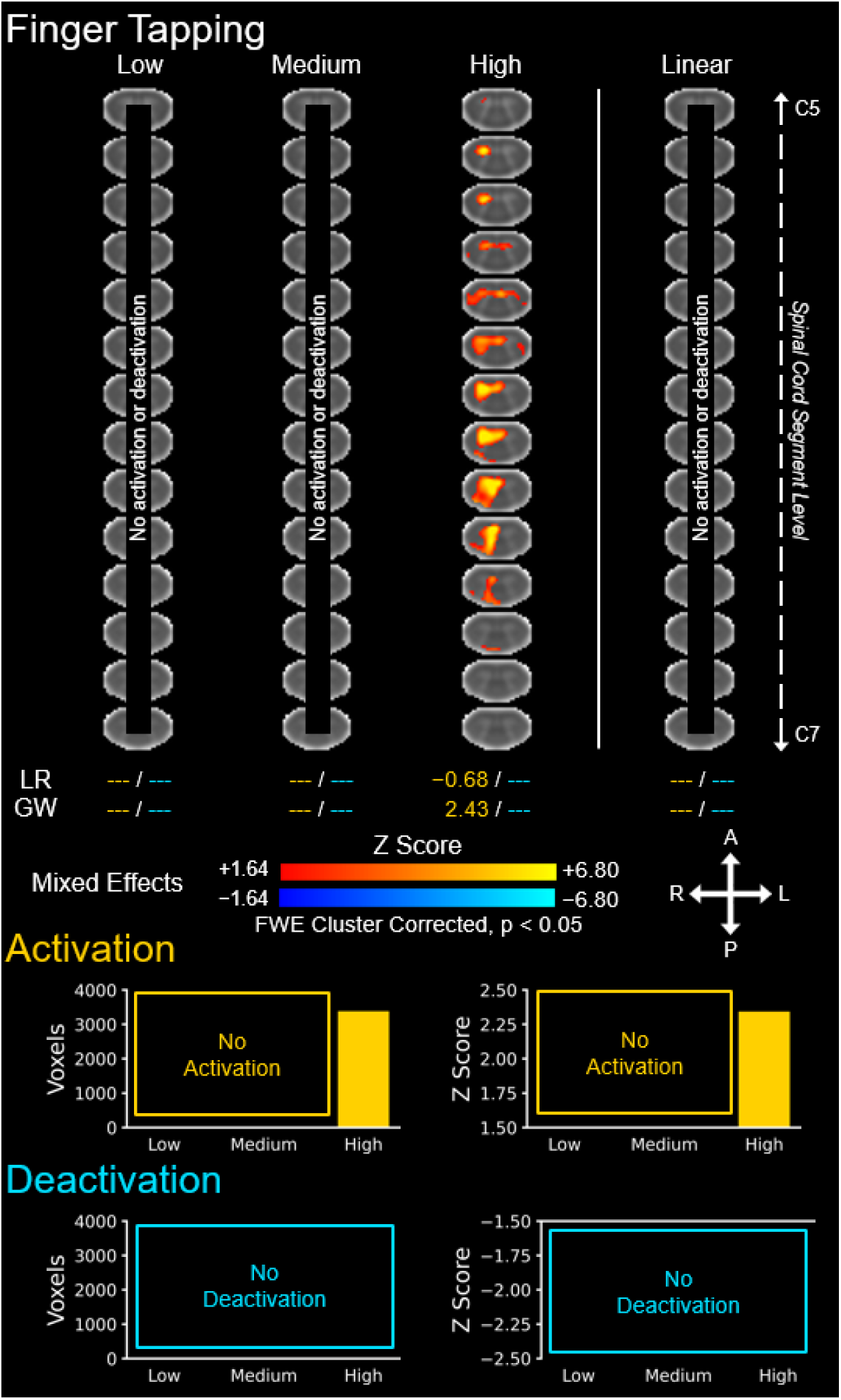
Group level spinal cord activity for the finger tapping task across the three task levels: low, medium, and high. Activations (i.e., positive signal change) are shown in red–yellow and deactivations are shown in blue–light blue (negative signal change). A linear contrast across the task levels was applied to map where the signal linearly increases and decreases across the task levels. The location of the activations and deactivations was assessed using the left-right (LR) index and gray matter-white matter (GW) ratio(--- = no activity, unable to calculate). The number of active voxels and the average Z score of the active voxels are shown to summarize the spatial extent and magnitude of the activations and deactivations across the three task levels. The number of active voxels and the average Z score of the active voxels are shown to summarize the spatial extent and magnitude of the activity across the three task levels. The activation maps were generated from a mixed effects analysis at the group level and were voxel-wise thresholded at a Z score > 1.64 with a family-wise error (FWE) cluster correction threshold of p < 0.05. The background image is the PAM50 T2*-weighted spinal cord template. Every 5th axial slice from the intersection of the subject level functional images is shown. A = anterior, P = posterior, L = left, R = right. --- = not applicable.

**Figure S6.**
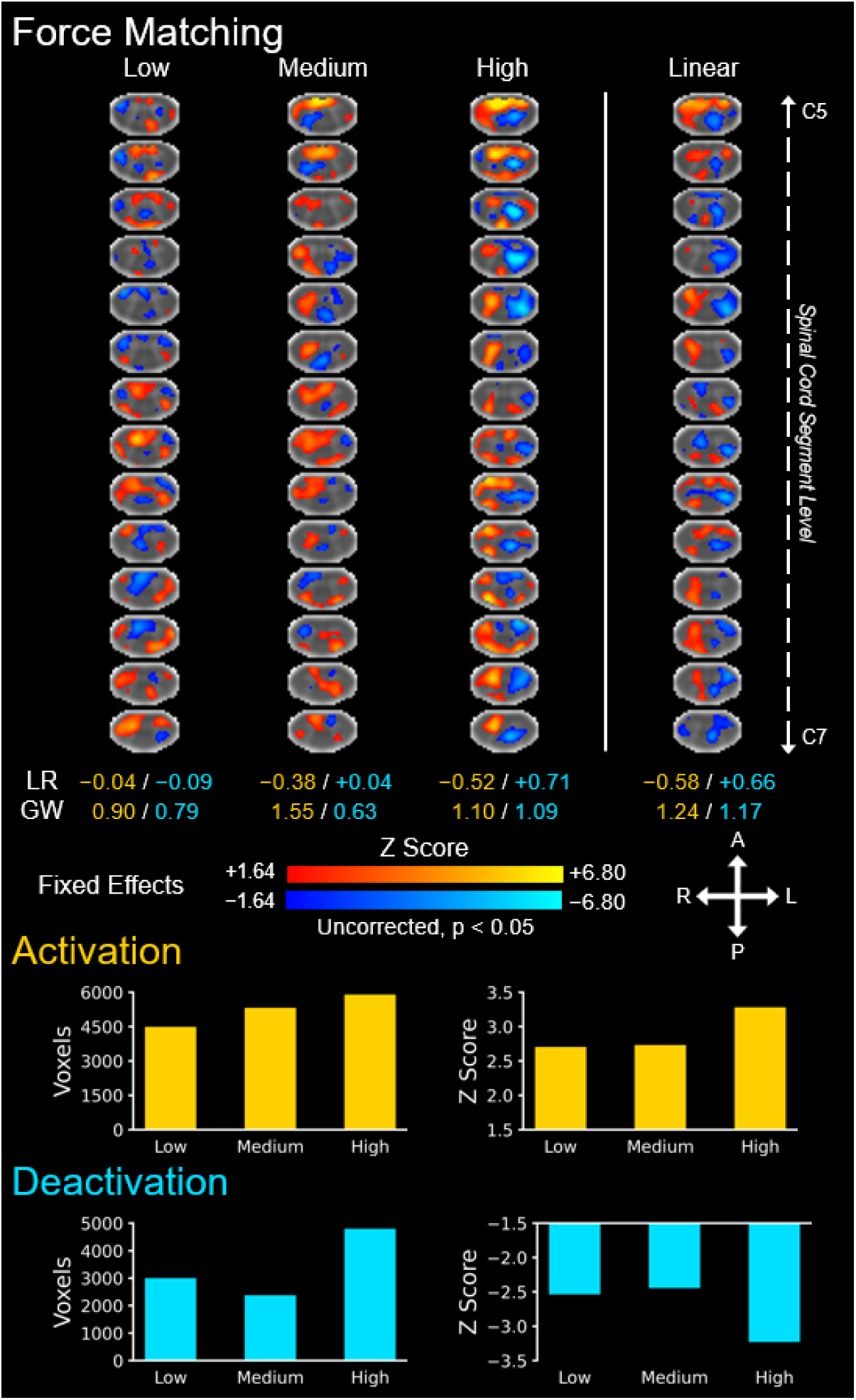
Group level spinal cord activity for the force matching task across the three task levels: low, medium, and high. Activations (i.e., positive signal change) are shown in red–yellow and deactivations are shown in blue–light blue (negative signal change). A linear contrast across the task levels was applied to map where the signal linearly increases and decreases across the task levels. The location of the activations and deactivations was assessed using the left-right (LR) index and gray matter-white matter (GW) ratio(--- = no activity, unable to calculate). The number of active voxels and the average Z score of the active voxels are shown to summarize the spatial extent and magnitude of the activations and deactivations across the three task levels. The number of active voxels and the average Z score of the active voxels are shown to summarize the spatial extent and magnitude of the activity across the three task levels. The activation maps were generated from a fixed effects analysis at the group level and were voxel-wise thresholded at a Z score > 1.64 without family-wise error correction (uncorrected). The background image is the PAM50 T2*-weighted spinal cord template. Every 5th axial slice from the intersection of the subject level functional images is shown. A = anterior, P = posterior, L = left, R = right.

**Figure S7.**
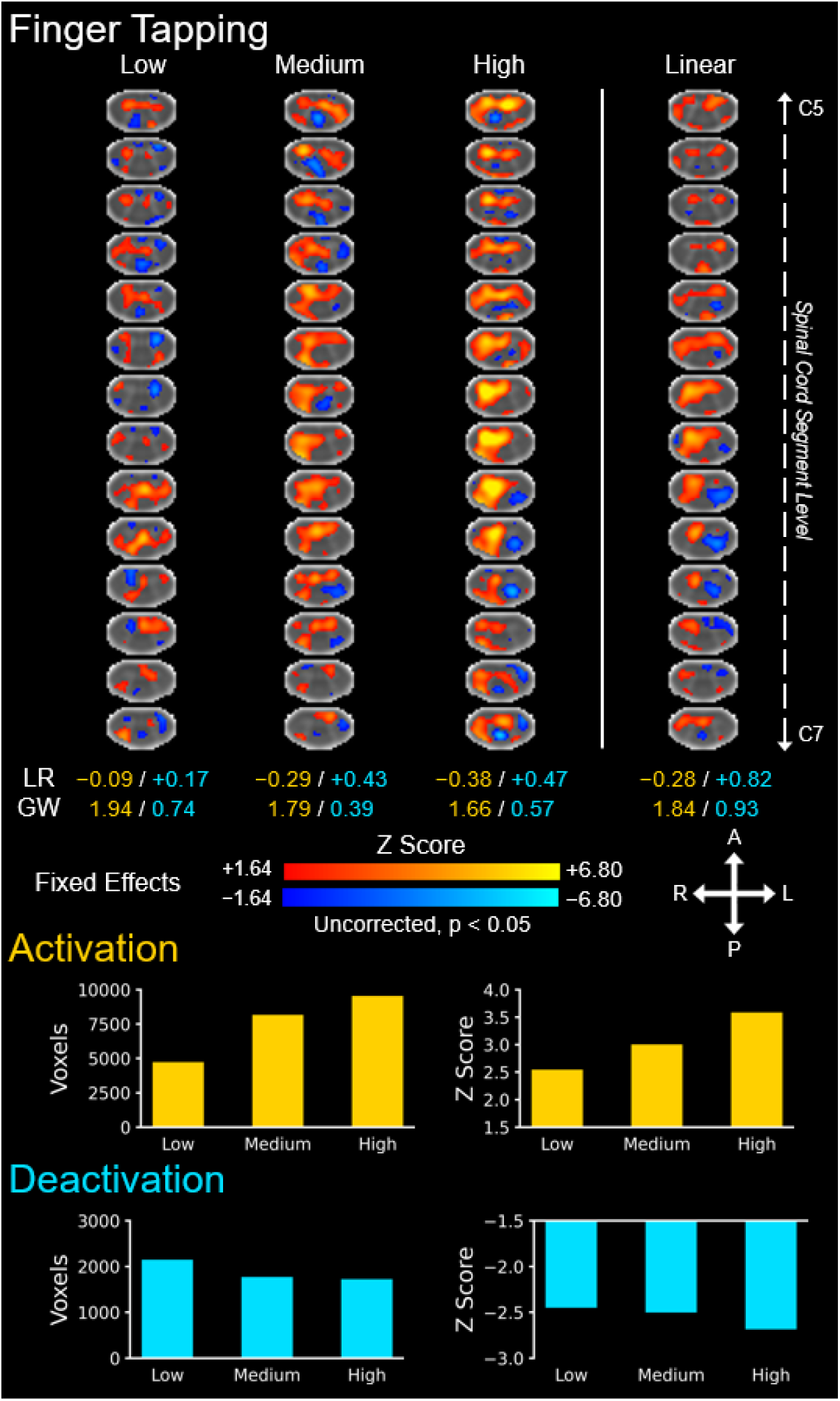
Group level spinal cord activity for the finger tapping task across the three task levels: low, medium, and high. Activations (i.e., positive signal change) are shown in red–yellow and deactivations are shown in blue–light blue (negative signal change). A linear contrast across the task levels was applied to map where the signal linearly increases and decreases across the task levels. The location of the activations and deactivations was assessed using the left-right (LR) index and gray matter-white matter (GW) ratio(--- = no activity, unable to calculate). The number of active voxels and the average Z score of the active voxels are shown to summarize the spatial extent and magnitude of the activations and deactivations across the three task levels. The number of active voxels and the average Z score of the active voxels are shown to summarize the spatial extent and magnitude of the activity across the three task levels. The activation maps were generated from a fixed effects analysis at the group level and were voxel-wise thresholded at a Z score > 1.64 without family-wise error correction (uncorrected). The background image is the PAM50 T2*-weighted spinal cord template. Every 5th axial slice from the intersection of the subject level functional images is shown. A = anterior, P = posterior, L = left, R = right.

**Figure S8.**
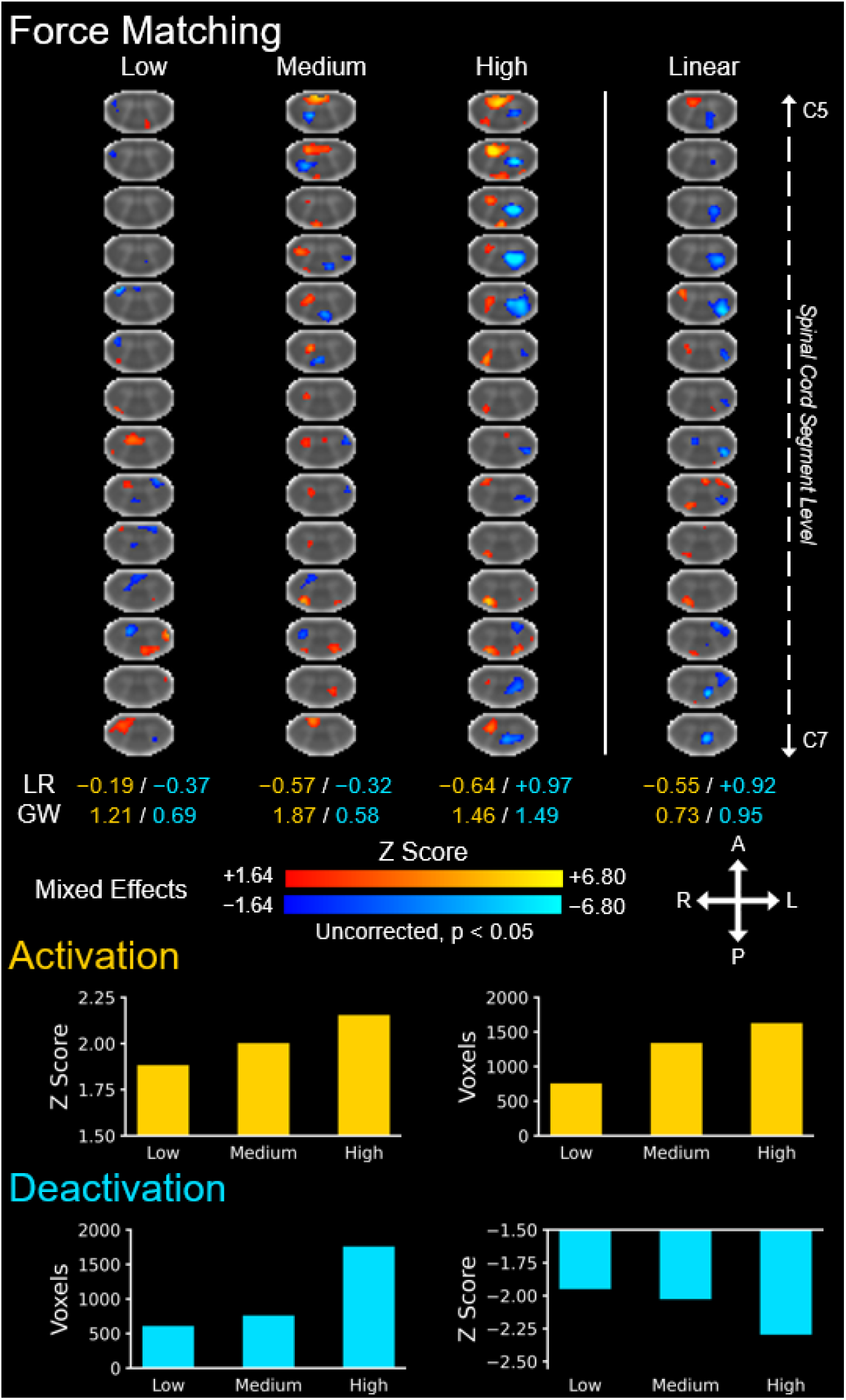
Group level spinal cord activity for the force matching task across the three task levels: low, medium, and high. Activations (i.e., positive signal change) are shown in red–yellow and deactivations are shown in blue–light blue (negative signal change). A linear contrast across the task levels was applied to map where the signal linearly increases and decreases across the task levels. The location of the activations and deactivations was assessed using the left-right (LR) index and gray matter-white matter (GW) ratio(--- = no activity, unable to calculate). The number of active voxels and the average Z score of the active voxels are shown to summarize the spatial extent and magnitude of the activations and deactivations across the three task levels. The number of active voxels and the average Z score of the active voxels are shown to summarize the spatial extent and magnitude of the activity across the three task levels. The activation maps were generated from a mixed effects analysis at the group level and were voxel-wise thresholded at a Z score > 1.64 without family-wise error correction (uncorrected). The background image is the PAM50 T2*-weighted spinal cord template. Every 5th axial slice from the intersection of the subject level functional images is shown. A = anterior, P = posterior, L = left, R = right.

**Figure S9.**
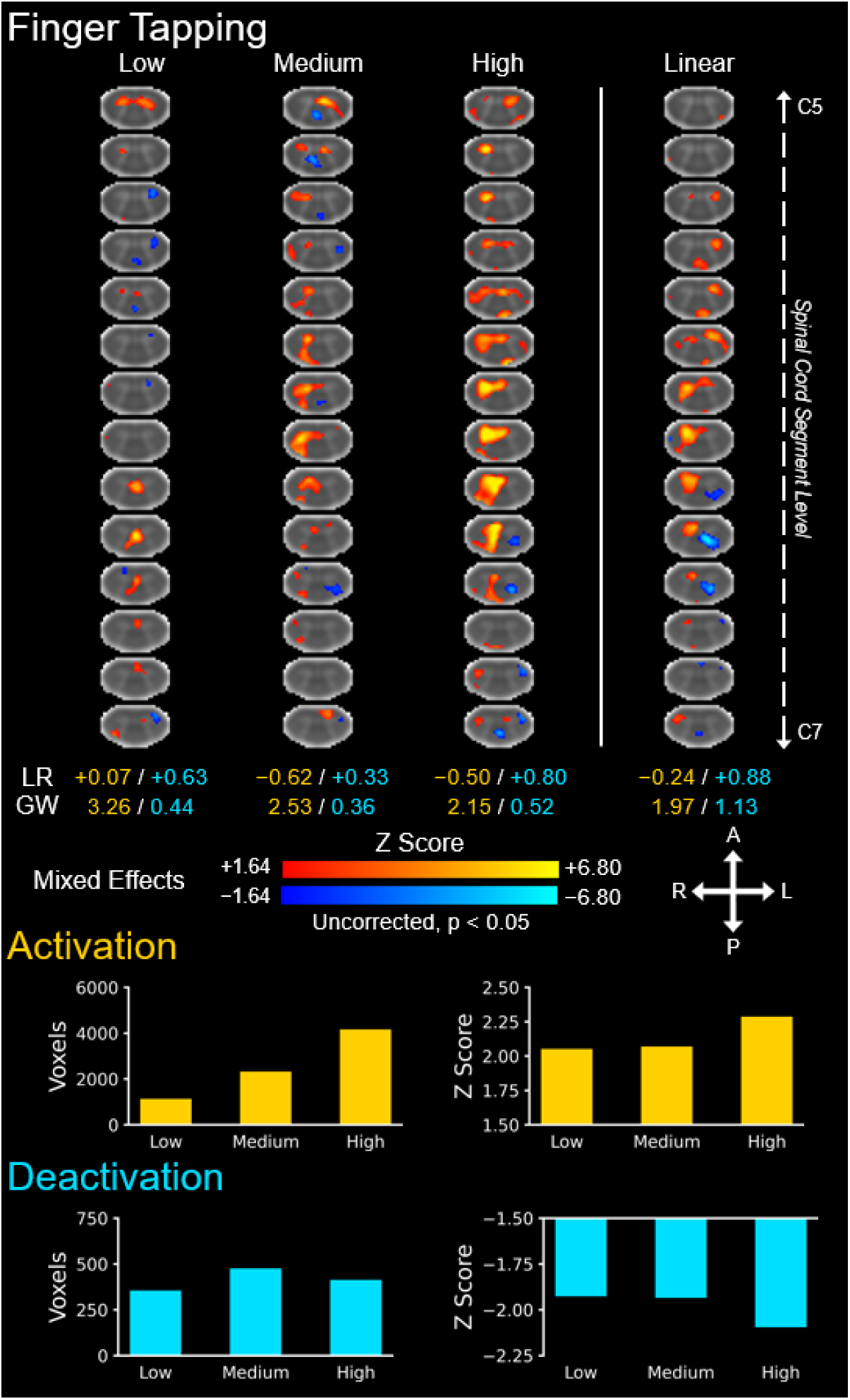
Group level spinal cord activity for the finger tapping task across the three task levels: low, medium, and high. Activations (i.e., positive signal change) are shown in red–yellow and deactivations are shown in blue–light blue (negative signal change). A linear contrast across the task levels was applied to map where the signal linearly increases and decreases across the task levels. The location of the activations and deactivations was assessed using the left-right (LR) index and gray matter-white matter (GW) ratio(--- = no activity, unable to calculate). The number of active voxels and the average Z score of the active voxels are shown to summarize the spatial extent and magnitude of the activations and deactivations across the three task levels. The number of active voxels and the average Z score of the active voxels are shown to summarize the spatial extent and magnitude of the activity across the three task levels. The activation maps were generated from a mixed effects analysis at the group level and were voxel-wise thresholded at a Z score > 1.64 without family-wise error correction (uncorrected). The background image is the PAM50 T2*-weighted spinal cord template. Every 5th axial slice from the intersection of the subject level functional images is shown. A = anterior, P = posterior, L = left, R = right.

## Notes

### Competing Interest Statement

The authors have declared no competing interest.

## References

Andersson, J. L. R., Jenkinson, M., & Smith, S. (n.d.). Non-linear registration aka Spatial normalisation FMRIB Technial Report TR07JA2. Retrieved July 26, 2024, from https://www.fmrib.ox.ac.uk/datasets/techrep/tr07ja2/tr07ja2.pdf

Avants, B. B., Tustison, N. J., Wu, J., Cook, P. A., & Gee, J. C. (2011). An open source multivariate framework for n-tissue segmentation with evaluation on public data. Neuroinformatics, 9(4), 381–400. 10.1007/s12021-011-9109-y

Beaton, D. E., Wright, J. G., Katz, J. N., & Upper Extremity Collaborative Group. (2005). Development of the QuickDASH: comparison of three item-reduction approaches. The Journal of Bone and Joint Surgery. American Volume, 87(5), 1038–1046. 10.2106/JBJS.D.02060

Bédard, S., Karthik, E. N., Tsagkas, C., Pravatà, E., Granziera, C., Smith, A., Weber, K. A., II, & Cohen-Adad, J. (2025). Towards contrast-agnostic soft segmentation of the spinal cord. Medical Image Analysis, 103473, 103473. 10.1016/j.media.2025.103473

Binkofski, F., Buccino, G., Posse, S., Seitz, R. J., Rizzolatti, G., & Freund, H. (1999). A fronto-parietal circuit for object manipulation in man: evidence from an fMRI-study: Fronto-parietal circuit for object manipulation. The European Journal of Neuroscience, 11(9), 3276–3286. 10.1046/j.1460-9568.1999.00753.x

Bonnard, M., Galléa, C., De Graaf, J. B., & Pailhous, J. (2007). Corticospinal control of the thumb-index grip depends on precision of force control: a transcranial magnetic stimulation and functional magnetic resonance imagery study in humans: Corticospinal control of precision grip. The European Journal of Neuroscience, 25(3), 872–880. 10.1111/j.1460-9568.2007.05320.x

Bortoff, G. A., & Strick, P. L. (1993). Corticospinal terminations in two new-world primates: further evidence that corticomotoneuronal connections provide part of the neural substrate for manual dexterity. The Journal of Neuroscience: The Official Journal of the Society for Neuroscience, 13(12), 5105–5118. 10.1523/jneurosci.13-12-05105.1993

Braaß, H., Feldheim, J., Chu, Y., Tinnermann, A., Finsterbusch, J., Büchel, C., Schulz, R., & Gerloff, C. (2023). Association between activity in the ventral premotor cortex and spinal cord activation during force generation-A combined cortico-spinal fMRI study. Human Brain Mapping, 44(18), 6471–6483. 10.1002/hbm.26523

Brooks, J. C. W., Beckmann, C. F., Miller, K. L., Wise, R. G., Porro, C. A., Tracey, I., & Jenkinson, M. (2008). Physiological noise modelling for spinal functional magnetic resonance imaging studies. NeuroImage, 39(2), 680–692. 10.1016/j.neuroimage.2007.09.018

Cadotte, D. W., Cadotte, A., Cohen-Adad, J., Fleet, D., Livne, M., Wilson, J. R., Mikulis, D., Nugaeva, N., & Fehlings, M. G. (2015). Characterizing the Location of Spinal and Vertebral Levels in the Human Cervical Spinal Cord. AJNR. American Journal of Neuroradiology, 36(4), 803–810. 10.3174/ajnr.A4192

Castiello, U., & Begliomini, C. (2008). The cortical control of visually guided grasping. The Neuroscientist: A Review Journal Bringing Neurobiology, Neurology and Psychiatry, 14(2), 157–170. 10.1177/1073858407312080

Cohen-Adad, J., Alonso-Ortiz, E., Abramovic, M., Arneitz, C., Atcheson, N., Barlow, L., Barry, R. L., Barth, M., Battiston, M., Büchel, C., Budde, M., Callot, V., Combes, A. J. E., De Leener, B., Descoteaux, M., de Sousa, P. L., Dostál, M., Doyon, J., Dvorak, A., … Xu, J. (2021). Generic acquisition protocol for quantitative MRI of the spinal cord. Nature Protocols, 16(10), 4611–4632. 10.1038/s41596-021-00588-0

Cona, G., & Semenza, C. (2017). Supplementary motor area as key structure for domain-general sequence processing: A unified account. Neuroscience and Biobehavioral Reviews, 72, 28–42. 10.1016/j.neubiorev.2016.10.033

Cramer, S. C., Weisskoff, R. M., Schaechter, J. D., Nelles, G., Foley, M., Finklestein, S. P., & Rosen, B. R. (2002). Motor cortex activation is related to force of squeezing. Human Brain Mapping, 16(4), 197–205. 10.1002/hbm.10040

Dai, T. H., Liu, J. Z., Sahgal, V., Brown, R. W., & Yue, G. H. (2001). Relationship between muscle output and functional MRI-measured brain activation. Experimental Brain Research. Experimentelle Hirnforschung. Experimentation Cerebrale, 140(3), 290–300. 10.1007/s002210100815

Dalley, A. F., II, & Agur, A. (2023). Moore’s clinically oriented anatomy (9th ed.). Wolters Kluwer Health. https://www.libreriastudium.it/it_IT/products/moore-s-clinically-oriented-anatomy

Davare, M., Andres, M., Cosnard, G., Thonnard, J.-L., & Olivier, E. (2006). Dissociating the role of ventral and dorsal premotor cortex in precision grasping. The Journal of Neuroscience: The Official Journal of the Society for Neuroscience, 26(8), 2260–2268. 10.1523/JNEUROSCI.3386-05.2006

De Leener, B., Fonov, V. S., Collins, D. L., Callot, V., Stikov, N., & Cohen-Adad, J. (2018). PAM50: Unbiased multimodal template of the brainstem and spinal cord aligned with the ICBM152 space. NeuroImage, 165, 170–179. 10.1016/j.neuroimage.2017.10.041

De Leener, B., Kadoury, S., & Cohen-Adad, J. (2014). Robust, accurate and fast automatic segmentation of the spinal cord. NeuroImage, 98, 528–536. 10.1016/j.neuroimage.2014.04.051

De Leener, B., Lévy, S., Dupont, S. M., Fonov, V. S., Stikov, N., Louis Collins, D., Callot, V., & Cohen-Adad, J. (2017). SCT: Spinal Cord Toolbox, an open-source software for processing spinal cord MRI data. NeuroImage, 145(Pt A), 24–43. 10.1016/j.neuroimage.2016.10.009

Ferbert, A., Priori, A., Rothwell, J. C., Day, B. L., Colebatch, J. G., & Marsden, C. D. (1992). Interhemispheric inhibition of the human motor cortex. The Journal of Physiology, 453(1), 525–546. 10.1113/jphysiol.1992.sp019243

Filippi, M., Sarasso, E., & Agosta, F. (2019). Resting-state functional MRI in parkinsonian syndromes: Resting-state fMRI in parkinsonian syndromes. Movement Disorders Clinical Practice, 6(2), 104–117. 10.1002/mdc3.12730

Finsterbusch, J., Eippert, F., & Büchel, C. (2012). Single, slice-specific z-shim gradient pulses improve T2*-weighted imaging of the spinal cord. NeuroImage, 59(3), 2307–2315. 10.1016/j.neuroimage.2011.09.038

Fratini, M., Moraschi, M., Maraviglia, B., & Giove, F. (2014). On the impact of physiological noise in spinal cord functional MRI: Physiological Noise in Spinal Cord fMRI. Journal of Magnetic Resonance Imaging: JMRI, 40(4), 770–777. 10.1002/jmri.24467

Galléa, C., Graaf, J. B. de, Pailhous, J., & Bonnard, M. (2008). Error processing during online motor control depends on the response accuracy. Behavioural Brain Research, 193(1), 117–125. 10.1016/j.bbr.2008.05.014

Giove, F., Garreffa, G., Giulietti, G., Mangia, S., Colonnese, C., & Maraviglia, B. (2004). Issues about the fMRI of the human spinal cord. Magnetic Resonance Imaging, 22(10), 1505–1516. 10.1016/j.mri.2004.10.015

Giulietti, G., Giove, F., Garreffa, G., Colonnese, C., Mangia, S., & Maraviglia, B. (2008). Characterization of the functional response in the human spinal cord: Impulse-response function and linearity. NeuroImage, 42(2), 626–634. 10.1016/j.neuroimage.2008.05.006

Grefkes, C., & Fink, G. R. (2011). Reorganization of cerebral networks after stroke: new insights from neuroimaging with connectivity approaches. Brain: A Journal of Neurology, 134(Pt 5), 1264–1276. 10.1093/brain/awr033

Hanajima, R., Ugawa, Y., Machii, K., Mochizuki, H., Terao, Y., Enomoto, H., Furubayashi, T., Shiio, Y., Uesugi, H., & Kanazawa, I. (2001). Interhemispheric facilitation of the hand motor area in humans. The Journal of Physiology, 531(Pt 3), 849–859. 10.1111/j.1469-7793.2001.0849h.x

Hemmerling, K. J., Hoggarth, M. A., Sandhu, M. S., Parrish, T. B., & Bright, M. G. (2023). Spatial distribution of hand-grasp motor task activity in spinal cord functional magnetic resonance imaging. Human Brain Mapping. 10.1002/hbm.26458

Henneman, E. (1957). Relation between size of neurons and their susceptibility to discharge. Science (New York, N.Y.), 126(3287), 1345–1347. 10.1126/science.126.3287.1345

Hernandez-Charpak, S. D., Kinany, N., Ricchi, I., Schlienger, R., Mattera, L., Martuzzi, R., Nazarian, B., Demesmaeker, R., Rowald, A., Kavounoudias, A., Bloch, J., Courtine, G., & Van De Ville, D. (2025). Towards personalized mapping through lumbosacral spinal cord task fMRI. Imaging Neuroscience, 3. 10.1162/imag_a_00455

Hoshi, E., & Tanji, J. (2007). Distinctions between dorsal and ventral premotor areas: anatomical connectivity and functional properties. Current Opinion in Neurobiology, 17(2), 234–242. 10.1016/j.conb.2007.02.003

Hou, B. L., Bhatia, S., & Carpenter, J. S. (2016). Quantitative comparisons on hand motor functional areas determined by resting state and task BOLD fMRI and anatomical MRI for pre-surgical planning of patients with brain tumors. NeuroImage. Clinical, 11, 378–387. 10.1016/j.nicl.2016.03.003

Islam, H., Law, C. S. W., Weber, K. A., Mackey, S. C., & Glover, G. H. (2019). Dynamic per slice shimming for simultaneous brain and spinal cord fMRI. Magnetic Resonance in Medicine: Official Journal of the Society of Magnetic Resonance in Medicine / Society of Magnetic Resonance in Medicine, 81(2), 825–838. 10.1002/mrm.27388

Jenkinson, M., Beckmann, C. F., Behrens, T. E. J., Woolrich, M. W., & Smith, S. M. (2012). FSL. NeuroImage, 62(2), 782–790. 10.1016/j.neuroimage.2011.09.015

Jenkinson, M., & Smith, S. (2001). A global optimisation method for robust affine registration of brain images. Medical Image Analysis, 5(2), 143–156. 10.1016/s1361-8415(01)00036-6

Johansen-Berg, H., Dawes, H., Guy, C., Smith, S. M., Wade, D. T., & Matthews, P. M. (2002). Correlation between motor improvements and altered fMRI activity after rehabilitative therapy. Brain: A Journal of Neurology, 125(Pt 12), 2731–2742. 10.1093/brain/awf282

Kaat, A. J., Buckenmaier, C. T., 3rd, Cook, K. F., Rothrock, N. E., Schalet, B. D., Gershon, R. C., & Vrahas, M. S. (2019). The expansion and validation of a new upper extremity item bank for the Patient-Reported Outcomes Measurement Information System® (PROMIS). Journal of Patient-Reported Outcomes, 3(1), 69. 10.1186/s41687-019-0158-6

Kaptan, M., Pfyffer, D., Konstantopoulos, C. G., Law, C. S. W., Weber, K. A., Ii, Glover, G. H., & Mackey, S. (2024). Recent developments and future avenues for human corticospinal neuroimaging. Frontiers in Human Neuroscience, 18, 1339881. 10.3389/fnhum.2024.1339881

Karadimas, S. K., Satkunendrarajah, K., Laliberte, A. M., Ringuette, D., Weisspapir, I., Li, L., Gosgnach, S., & Fehlings, M. G. (2020). Sensory cortical control of movement. Nature Neuroscience, 23(1), 75–84. 10.1038/s41593-019-0536-7

Khatibi, A., Vahdat, S., Lungu, O., Finsterbusch, J., Büchel, C., Cohen-Adad, J., Marchand-Pauvert, V., & Doyon, J. (2022). Brain-spinal cord interaction in long-term motor sequence learning in human: An fMRI study. NeuroImage, 253(119111), 119111. 10.1016/j.neuroimage.2022.119111

Kinany, N., Khatibi, A., Lungu, O., Finsterbusch, J., Büchel, C., Marchand-Pauvert, V., Van De Ville, D., Vahdat, S., & Doyon, J. (2023). Decoding cerebro-spinal signatures of human behavior: Application to motor sequence learning. NeuroImage, 275(120174), 120174. 10.1016/j.neuroimage.2023.120174

Klein, A., & Tourville, J. (2012). 101 labeled brain images and a consistent human cortical labeling protocol. Frontiers in Neuroscience, 6, 171. 10.3389/fnins.2012.00171

Leskinen, S., Singha, S., Mehta, N. H., Quelle, M., Shah, H. A., & D’Amico, R. S. (2024). Applications of functional magnetic resonance imaging to the study of functional connectivity and activation in neurological disease: A scoping review of the literature. World Neurosurgery, 189, 185–192. 10.1016/j.wneu.2024.06.003

Madi, S., Flanders, A., Vinitski, S., Herbison, G., & Nissanov, J. (2001). Functional MR imaging of the human cervical spinal cord. AJNR. American Journal of Neuroradiology, 22(9), 1768–1774. https://www.ajnr.org/content/22/9/1768?ijkey=06077a67b939271acb08f271f1a442402a7581d5&keytype2=tf_ipsecsha#sec-8

Maehara, M., Ikeda, K., Kurokawa, H., Ohmura, N., Ikeda, S., Hirokawa, Y., Maehara, S., Utsunomiya, K., Tanigawa, N., & Sawada, S. (2014). Diffusion-weighted echo-planar imaging of the head and neck using 3-T MRI: Investigation into the usefulness of liquid perfluorocarbon pads and choice of optimal fat suppression method. Magnetic Resonance Imaging, 32(5), 440–445. 10.1016/j.mri.2014.01.011

Maieron, M., Iannetti, G. D., Bodurka, J., Tracey, I., Bandettini, P. A., & Porro, C. A. (2007). Functional responses in the human spinal cord during willed motor actions: evidence for side- and rate-dependent activity. The Journal of Neuroscience: The Official Journal of the Society for Neuroscience, 27(15), 4182–4190. 10.1523/JNEUROSCI.3910-06.2007

Mayka, M. A., Corcos, D. M., Leurgans, S. E., & Vaillancourt, D. E. (2006). Three-dimensional locations and boundaries of motor and premotor cortices as defined by functional brain imaging: a meta-analysis. NeuroImage, 31(4), 1453–1474. 10.1016/j.neuroimage.2006.02.004

Morecraft, R. J., Ge, J., Stilwell-Morecraft, K. S., Lemon, R. N., Ganguly, K., & Darling, W. G. (2023). Terminal organization of the corticospinal projection from the arm/hand region of the rostral primary motor cortex (M1r or old M1) to the cervical enlargement (C5-T1) in rhesus monkey. The Journal of Comparative Neurology, 531(18), 1996–2018. 10.1002/cne.25557

Nachev, P., Kennard, C., & Husain, M. (2008). Functional role of the supplementary and pre-supplementary motor areas. Nature Reviews. Neuroscience, 9(11), 856–869. 10.1038/nrn2478

Natali, A. L., Reddy, V., & Bordoni, B. (2024). Neuroanatomy, corticospinal cord tract. In StatPearls. StatPearls Publishing. https://www.ncbi.nlm.nih.gov/books/NBK535423/#:~:text=The%20corticospinal%20tract%2C%20AKA%2C%20the,movement%20of%20the%20distal%20extremities.

Ng, M.-C., Wu, E. X., Lau, H.-F., Hu, Y., Lam, E. Y., & Luk, K. D. (2008). Cervical spinal cord BOLD fMRI study: modulation of functional activation by dexterity of dominant and non-dominant hands. NeuroImage, 39(2), 825–831. 10.1016/j.neuroimage.2007.09.026

Reuben, D. B., Magasi, S., McCreath, H. E., Bohannon, R. W., Wang, Y.-C., Bubela, D. J., Rymer, W. Z., Beaumont, J., Rine, R. M., Lai, J.-S., & Gershon, R. C. (2013). Motor assessment using the NIH Toolbox. Neurology, 80(11 Suppl 3), S65–S75. 10.1212/WNL.0b013e3182872e01

rpy2: Python-R bridge. (n.d.). Retrieved August 13, 2024, from https://rpy2.github.io/

Steuer, I., & Guertin, P. A. (2019). Central pattern generators in the brainstem and spinal cord: an overview of basic principles, similarities and differences. Reviews in the Neurosciences, 30(2), 107–164. 10.1515/revneuro-2017-0102

Stroman, P. W., & Ryner, L. N. (2001). Functional MRI of motor and sensory activation in the human spinal cord. Magnetic Resonance Imaging, 19(1), 27–32. 10.1016/s0730-725x(01)00226-0

Tanji, J. (2001). Sequential organization of multiple movements: involvement of cortical motor areas. Annual Review of Neuroscience, 24(1), 631–651. 10.1146/annurev.neuro.24.1.631

Ueno, M., Nakamura, Y., Li, J., Gu, Z., Niehaus, J., Maezawa, M., Crone, S. A., Goulding, M., Baccei, M. L., & Yoshida, Y. (2018). Corticospinal circuits from the sensory and motor cortices differentially regulate skilled movements through distinct spinal interneurons. Cell Reports, 23(5), 1286–1300.e7. 10.1016/j.celrep.2018.03.137

Ullmann, E., Pelletier Paquette, J. F., Thong, W. E., & Cohen-Adad, J. (2014). Automatic labeling of vertebral levels using a robust template-based approach. International Journal of Biomedical Imaging, 2014, 719520. 10.1155/2014/719520

Vahdat, S., Lungu, O., Cohen-Adad, J., Marchand-Pauvert, V., Benali, H., & Doyon, J. (2015). Simultaneous Brain–Cervical Cord fMRI Reveals Intrinsic Spinal Cord Plasticity during Motor Sequence Learning. PLoS Biology, 13(6), e1002186. 10.1371/journal.pbio.1002186

Valošek, J., Mathieu, T., Schlienger, R., Kowalczyk, O. S., & Cohen-Adad, J. (2024). Automatic segmentation of the spinal cord nerve rootlets. Imaging Neuroscience, 2, 1–14. 10.1162/imag_a_00218

Veale, J. F. (2014). Edinburgh Handedness Inventory - Short Form: a revised version based on confirmatory factor analysis. Laterality, 19(2), 164–177. 10.1080/1357650X.2013.783045

Weber, K. A., 2nd, Chen, Y., Wang, X., Kahnt, T., & Parrish, T. B. (2016). Lateralization of cervical spinal cord activity during an isometric upper extremity motor task with functional magnetic resonance imaging. NeuroImage, 125, 233–243. 10.1016/j.neuroimage.2015.10.014

Weber, K. A., Chen, Y., Wang, X., & Parrish, T. B. (2014). Choice of motion correction method affects spinal cord fMRI results. 20th Annual Meeting of the Organization for Human Brain Mapping.

Witt, S. T., Laird, A. R., & Meyerand, M. E. (2008). Functional neuroimaging correlates of finger-tapping task variations: an ALE meta-analysis. NeuroImage, 42(1), 343–356. 10.1016/j.neuroimage.2008.04.025

Wong, A. L., Haith, A. M., & Krakauer, J. W. (2015). Motor planning. The Neuroscientist: A Review Journal Bringing Neurobiology, Neurology and Psychiatry, 21(4), 385–398. 10.1177/1073858414541484

Xie, C.-H., Kong, K.-M., Guan, J.-T., Chen, Y.-X., He, J.-K., Qi, W.-L., Wang, X.-J., Shen, Z.-W., & Wu, R.-H. (2009). SSFSE sequence functional MRI of the human cervical spinal cord with complex finger tapping. European Journal of Radiology, 70(1), 1–6. 10.1016/j.ejrad.2008.01.003

Yarkoni, T., Poldrack, R. A., Nichols, T. E., Van Essen, D. C., & Wager, T. D. (2011). Large-scale automated synthesis of human functional neuroimaging data. Nature Methods, 8(8), 665–670. 10.1038/nmeth.1635

Zehr, E. P., Carroll, T. J., Chua, R., Collins, D. F., Frigon, A., Haridas, C., Hundza, S. R., & Thompson, A. K. (2004). Possible contributions of CPG activity to the control of rhythmic human arm movement. Canadian Journal of Physiology and Pharmacology, 82(8-9), 556–568. 10.1139/y04-056

Zhang, Y., Brady, M., & Smith, S. (2001). Segmentation of brain MR images through a hidden Markov random field model and the expectation-maximization algorithm. IEEE Transactions on Medical Imaging, 20(1), 45–57. 10.1109/42.906424

